# A *Plasmodium knowlesi* A1-H.1 transcriptome time course focusing on the late asexual blood stages

**DOI:** 10.64898/2026.02.09.704036

**Authors:** Katlijn De Meulenaere, Dalia Díaz-Delgado, Pieter Monsieurs, Erin Sauve, Alfred Cortés, Ellen Knuepfer, Anna Rosanas-Urgell

## Abstract

**Background:** The zoonotic parasite *Plasmodium knowlesi* is closely related to *Plasmodium vivax*, the second leading cause of human malaria. *P. knowlesi* line A1-H.1 can be maintained in human erythrocytes and is an experimental model for *P. vivax* biology. We present the first transcriptome time-course of the intraerythrocytic developmental cycle (IDC) of *P. knowlesi* A1-H.1 grown in human normocytes.

**Results:** Bulk RNA-sequencing was performed at five IDC stages using tightly synchronised cultures, enabling identification of constitutively expressed genes and stage-specific biomarkers, and investigation of temporal expression patterns of multigene families. Comparative analysis with *P. vivax* orthologues revealed strong genome-wide conservation of temporal expression. Analysis of invasion-associated genes and ApiAP2 transcription factors identified genes with conserved expression patterns for which *P. knowlesi* is likely a good model to study the *P. vivax* gene. To support comparative analyses, we developed an open-access interactive webtool to explore and visualise *P. knowlesi* – *P. vivax* orthologue expression across the IDC.

**Conclusions:** This time-course dataset provides a reference transcriptome framework for *P. knowlesi* A1-H.1 and a resource for comparative *Plasmodium* biology. The interactive webtool facilitates rapid identification of candidate genes with conserved temporal expression, streamlining functional studies in which *P. knowlesi* A1-H.1 serves as a model for *P. vivax*.

## Background

The simian parasite *Plasmodium knowlesi* is phylogenetically closely related to *Plasmodium vivax* (1–3) and, unlike *P. vivax*, can be long-term cultured *in vitro*. In nature, *P. knowlesi* primarily infects long- and pig-tailed macaques in Southeast Asia, but also causes zoonotic human infections, predominantly described from Malaysia (4). Thus, *P. knowlesi* can infect and replicate in both human and macaque erythrocytes. Similar to *Plasmodium falciparum* but in contrast to *P. vivax, P. knowlesi* can invade both normocytes and reticulocytes, although it has a preference for reticulocytes (5). While *P. falciparum* belongs to the subgenus *Laverania*, both *P. knowlesi* and *P. vivax* are members of the subgenus *Plasmodium* and are therefore more closely related. This is reflected in the high degree of genomic synteny between *P. knowlesi* and *P. vivax* (2, 6), supporting the suitability of *P. knowlesi* laboratory strains as an experimental model to investigate *P. vivax* gene functions (7).

*P. knowlesi* strain H was isolated from a clinical human infection (8) and subsequently passaged through rhesus macaques. One resulting clone, Pk1(A+) (9), was sequenced to generate the first *P. knowlesi* reference genome ‘H’ (10). More recently, strain A1, derived from strain H, was gradually adapted to grow in human erythrocytes, while it maintained the capacity to grow in macaque erythrocytes. The resulting A1-H.1 line (‘Pk A1-H.1’) (11) is particularly suited to investigate *P. knowlesi* processes involving human red blood cells (RBCs), and eliminates the need for macaque RBCs, which are often difficult to procure. Pk A1-H.1 has been shown to be a valuable tool for drug and vaccine studies (12–14). Furthermore, the development of a CRISPR/Cas9 system for genetic manipulation of Pk A1-H.1 (15) allows targeted studies of *P. vivax* orthologues. These combined characteristics establish *P. knowlesi* line A1-H.1 as a powerful model for advancing *P. vivax* research.

*P. vivax* is the most wide-spread malaria-causing parasite and is responsible for the second highest number of cases after *P. falciparum* (16). However, it remains under-researched due to the lack of a continuous *in vitro* culturing system. Consequently, functional assays, genetic engineering and early-stage drug or vaccine testing is limited if at all possible (17, 18); yet these experiments are crucial to explore *P. vivax* biology, and to develop effective strategies for prevention, control and elimination. To investigate *P. vivax* gene function using *P. knowlesi* as a model, it is not only important to identify *P. knowlesi* orthologues of the *P. vivax* genes of interest, but also to ensure similar temporal transcriptional profiles between the orthologues, which increases the likelihood that they participate in related biological processes. Transcriptome datasets covering *P. knowlesi* stages across the intraerythrocytic developmental cycle (IDC) are essential for this purpose. The majority of genes in *Plasmodium* are expressed in a ‘just-in-time’ pattern during the IDC (19–22), where genes are usually sequentially activated at a specific time when their product is needed. Life cycle progression is mainly regulated at the transcriptional level by Apicomplexan apetala2 (AP2) domain-containing transcription factors (ApiAP2) (23–26).

For Pk A1-H.1, no bulk RNA-sequencing (RNA-seq) time course dataset is currently available. The only *P. knowlesi* transcriptome time course datasets available at the time of writing were generated from clone Pk1(A+) (21). In that study, Pk1(A+) parasites were obtained from infected rhesus macaques or *in vitro* cultures using macaque erythrocytes. The similarity of these transcriptome time courses with the transcriptome of Pk A1-H.1 grown in human RBCs is expected to be limited, especially for genes related to RBC tropism. Furthermore, the existing time course datasets were generated using microarray technology based on an older annotation of the *P. knowlesi* reference genome H. Hence, more recently annotated genes were not detected. Since then, two *P. knowlesi* A1-H.1 single-cell RNA-seq (scRNA-seq) datasets were published (27, 28). Both datasets contain all asexual IDC stages and provide valuable insights into transcription patterns across the asexual developmental cycle in the blood. While scRNA-seq excels at resolving cellular heterogeneity, its lower sequencing depth of individual genes compared to bulk RNA-seq reduces the sensitivity to detect genes with low expression levels and makes scRNA-seq less suitable for generating high-resolution temporal expression profiles (29, 30).

In this study, we generated the first transcriptome time course for the 27-hour asexual intraerythrocytic life cycle of *P. knowlesi* line A1-H.1, with a higher temporal resolution during trophozoite and schizont development. We collected *P. knowlesi* A1-H.1 RNA at five time points during the IDC: 5 hours post-invasion (hpi) rings, 14 hpi mid-trophozoites, 20 hpi late trophozoites, 24 hpi mid-schizonts and 27 hpi late schizonts, and generated transcriptomes using RNA-seq. We identified constitutively expressed genes and stage-specific biomarkers, and characterised the expression dynamics of multigene families. Temporal expression profiles were compared on a genome-wide scale with *P. vivax* orthologues, with a focus on invasion-associated genes and ApiAP2 transcription factors. To support these and broader comparative analyses, an interactive web tool was developed that allows visualisation of *P. knowlesi* and *P. vivax* gene expression patterns over the IDC, either for orthologous gene pairs or for individual genes analysed independently. We expect this tool to facilitate studying these human-infecting neglected *Plasmodium* parasites.

## Results

### Culturing of *P. knowlesi* A1-H.1 for transcriptome collection at 5 time points

*P. knowlesi* line A1-H.1 was grown in human erythrocytes to generate transcriptomes of 5 different life cycle stages during the IDC, *i.e.* 5, 14, 20, 24 and 27 hpi (**Fig. 1**). Cultures were tightly synchronised to a 1-hour age window. We monitored the progression of the parasite cultures through the IDC by microscopy of Giemsa-stained blood smears (**Fig. 1**). The 5 hpi and 14 hpi parasites showed ring and mid-trophozoite stages, respectively. The 20, 24 and 27 hpi late stages (late trophozoites, mid-schizonts and late schizonts) were purified on a Nycodenz bed prior to harvesting. At 27 hpi, we observed late segmenting schizonts with a fan-like shape and condensed hemozoin, along with free merozoites and ring stages (∼30% of all parasites before Nycodenz purification of schizonts), indicating that schizonts were mature and merozoites are egressing at this time point. The IDC length of Pk A1-H.1, reported as 27 hours (11), matched the observed progression of our parasite cultures.

**Figure 1.**
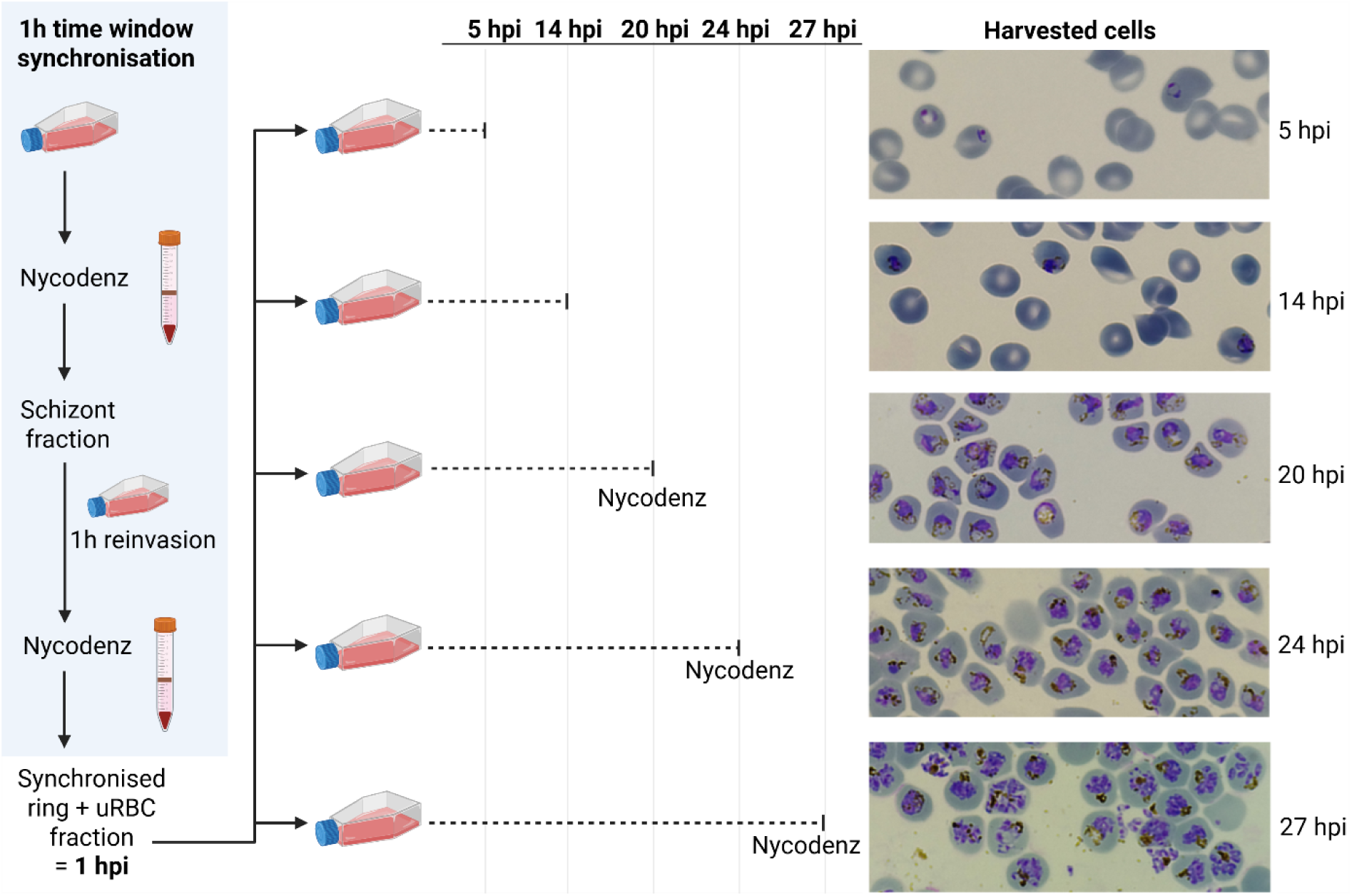
Overview of the transcriptome time course sample collection. *P. knowlesi* A1-H.1 parasites were synchronised to a 1-hour time window (blue field), the resulting fraction of rings and uninfected RBCs (uRBCs) was cultured and parasites harvested at 5, 14, 20, 24 and 27 hours post-invasion (hpi) for mRNA sequencing. The 20, 24 and 27 hpi samples were parasite-enriched on an isotonic 55% Nycodenz bed before harvest. The 20, 24 and 27 hpi flasks were prepared from the same synchronised culture (culture 3), while the 5 hpi and 14 hpi flasks were prepared from separately synchronised cultures (culture 1 and 2).

### The temporal transcriptional profile of the *P. knowlesi* A1-H.1 IDC

We compared gene expression across the 5 sampled time points to assess oscillations in transcript levels typical of the highly regulated *Plasmodium* IDC (19, 21, 31). A genome-wide heatmap, with genes ordered by their peak expression (**Fig. 2**), revealed a ‘just-in-time’ transcriptional programme, with a set of specific genes predominantly expressed at each time point. Temporal expression patterns of *P. knowlesi* genes of choice can be visualised and compared using our interactive web tool (https://interactive.itg.be/app/mal-pk-pv-expression-viewer).

**Figure 2.**
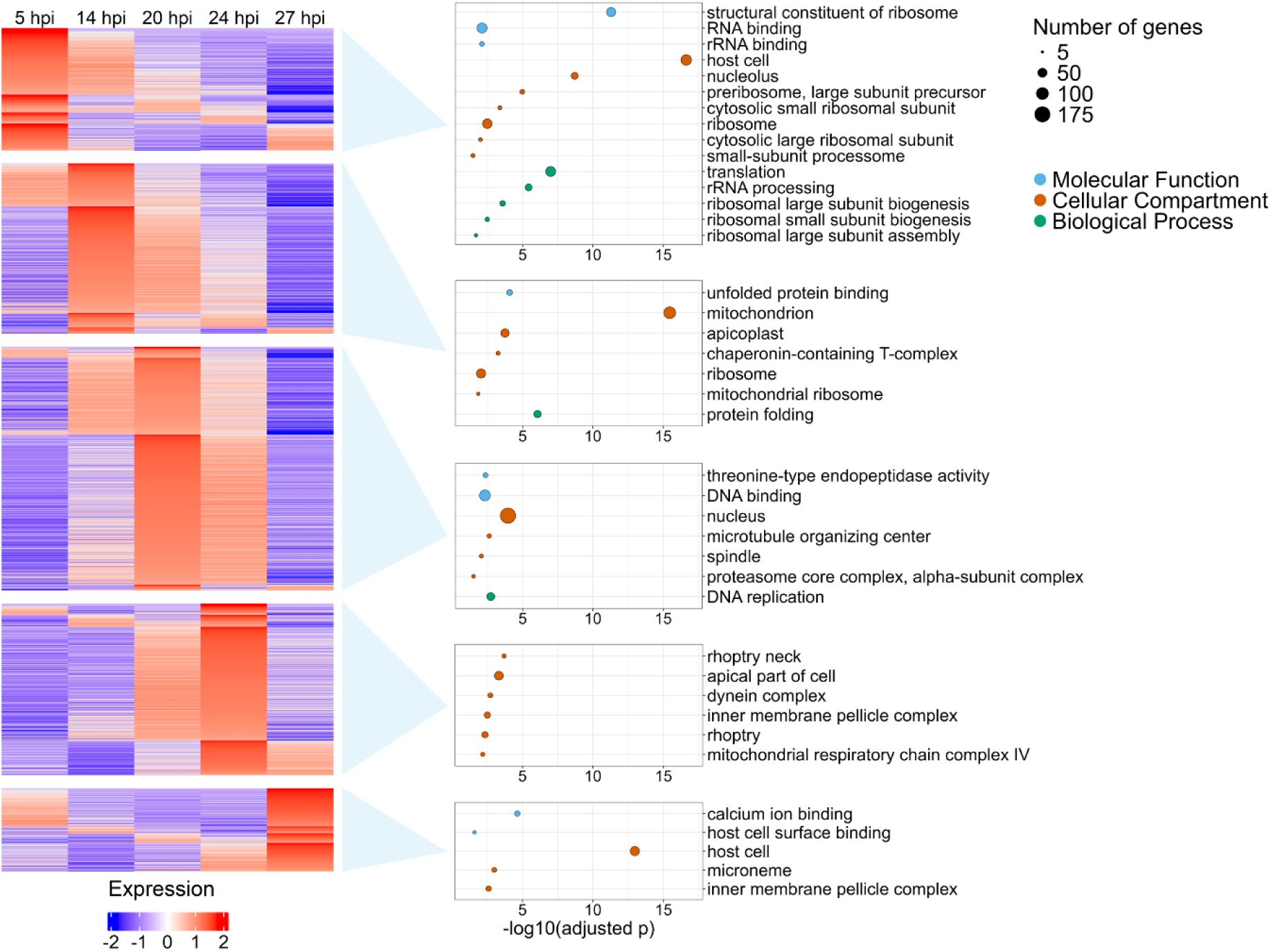
Temporal transcriptional pattern of *P. knowlesi* A1-H.1 over the IDC. Heatmap of normalised (TPM-normalised, log_2_-transformed and Z-scored) expression of all genes (rows) with sufficient expression and temporal variability (filtering criteria detailed in Methods), for the 5 collected time points (columns). Genes were grouped by their time point of peak expression, reflected by the heatmap blocks. Within each time point block, genes were additionally ordered by the time point at which they were second highest expressed. For each group of genes peaking at a specific time point (time point blocks), a gene ontology (GO) enrichment analysis was performed against a background of all *P. knowlesi* genes. Bubble plots display significantly enriched GO terms per time point (Bonferroni-adjusted p<0.05). The x-axis shows the negative logarithm of the p-value, with higher numbers indicating higher significance levels. Bubble size reflects the gene number; colours indicate the GO categories (Molecular Function, Cellular Compartment or Biological Process). If a category is not shown, no significant enrichment was detected.

Additionally, we investigated the genes with the highest expression levels at each time point to assess whether their biological functions correspond to the life cycle stage at which they are maximally expressed. We first selected the 10 most highly expressed nuclear-encoded genes (excluding apicoplast- and mitochondrion-encoded genes) per time point (**Supplementary Table S1**) and observed stage-specific differences. Among the top expressed genes were *Early Transcribed Membrane Protein* (*ETRAMP*) genes (5, 14, 27 hpi) and *Merozoite Surface Protein* (*MSP*) genes (20, 24 hpi) genes, as well as genes encoding for ribosomal proteins (5, 14 hpi) and histones (20, 24 hpi). Notably, the 20 hpi and 24 hpi time points showed overlap in their top expressed genes. At 27 hpi, parasites upregulate invasion machinery genes (actin I, IMC1c), together with parasitophorous vacuole membrane-associated genes (*ETRAMP*, *EXP1*) and exported genes, that may establish the new intracellular niche after RBC invasion.

We next performed a gene ontology (GO) enrichment analysis on gene sets grouped according to their peak expression time point (**Fig. 2, Additional File 2**). This revealed stage-specific biological functions: genes peaking at 5 hpi and 14 hpi are enriched in GO terms related to transcription and translation, genes peaking at the late trophozoite stage (20 hpi) are associated with DNA replication, and genes peaking at the mid- and late schizont stages (24 and 27 hpi) are enriched in invasion-related GO terms. Consistent with observations in *P. falciparum*, *P. knowlesi* A1-H.1 rhoptry proteins start being expressed during the mid-schizont stage (enrichment of ‘rhoptry’, ‘rhoptry neck’ GO terms), followed by expression of microneme proteins during the late schizonts (enrichment of GO term ‘microneme’) (19, 32). Of note, not all *P. knowlesi* genes have a GO annotation, and additional functional patterns may emerge as annotations improve (**Additional File 2**). Overall, *P. knowlesi* A1-H.1 exhibits the temporally regulated transcriptional and biological patterns characteristic of the *Plasmodium* IDC.

### Constitutively expressed genes and stage-specific biomarkers

Based on the transcriptional patterns over the IDC observed in this dataset, we extracted genes that could potentially be used for normalisation or stage identification in *P. knowlesi* research. Firstly, eight constitutively expressed genes with stable and high expression over the 5 time points were identified (**Fig. 3**). These genes are suitable for *e.g.* normalisation of reverse transcription quantitative PCR, or for their promoters to drive transgenic gene expression. As expected, these constitutively expressed genes mostly encode housekeeping proteins with general cellular functions, although one conserved protein of unknown function was also included.

**Figure 3.**
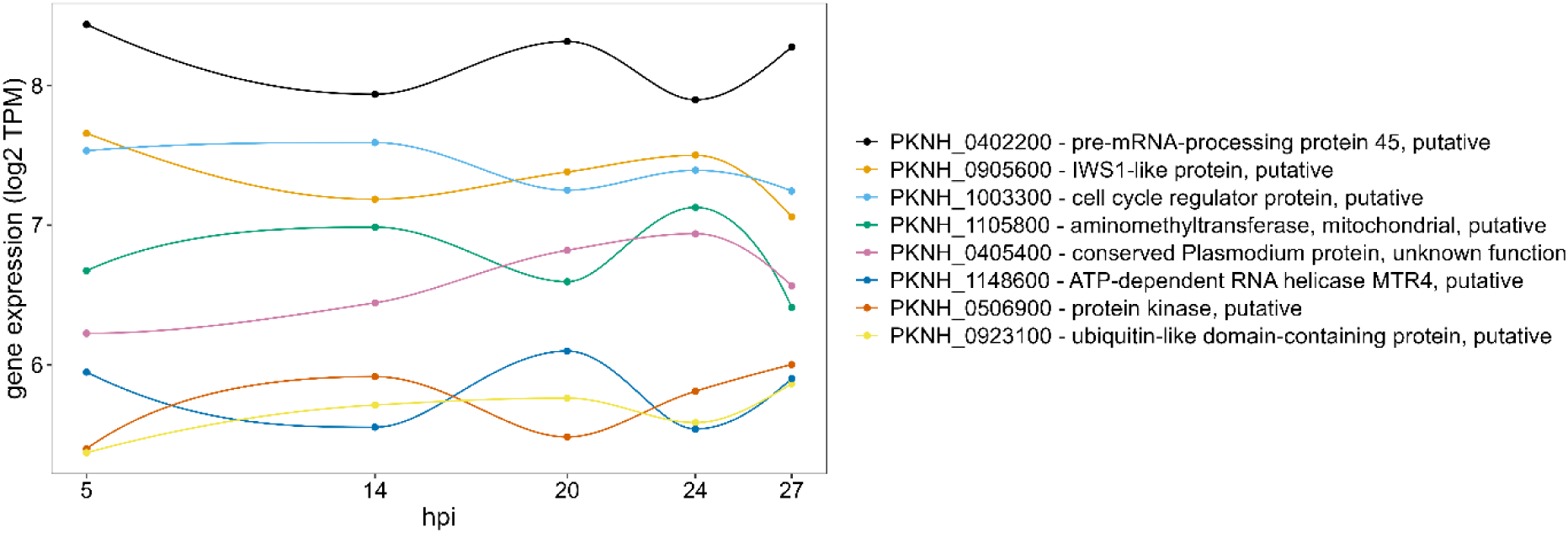
Expression levels of 8 constitutively expressed genes over the IDC. Normalised interpolated expression levels (TPM-normalised, log_2_-transformed) of 8 constitutively expressed genes are shown, which were selected based on their low transcriptional variation over the 5 discrete time points and high overall expression (selection criteria detailed in Methods). Dots reflect the 5 sampled time points; intermediate values were interpolated using PCHIP.

Secondly, for each time point, biomarker genes with stage-specific expression were identified based on differential expression (3-fold upregulated compared to every other time point), after which the 20 genes with the highest expression level at that time point were selected (**Fig. 4A, Additional File 3**). These biomarkers can for example be used for life cycle stage annotation (expressed in hpi) of *P. knowlesi* single-cell transcriptomes. The scRNA-sequenced parasite cells from these datasets are usually visualised in a UMAP plot where parasite cells with similar transcriptomic profiles are close to each other, revealing a cyclic organisation that reflects progression of cells through the IDC. Parasite cells on these UMAP plots require annotation with stage-specific markers for which the precise time window of expression is known, in order to assign each cell to its corresponding life cycle stage, assess the quality of the clustering underlying the UMAP, estimate gene expression timing, and compare different *P. knowlesi* single-cell datasets. In the existing *P. knowlesi* single-cell datasets, time point and life stage annotation is done based on microarray data from *P. berghei* (27, 28). Because single-cell annotations are ideally based on species-matched datasets, and because bulk RNA-seq provides high-quality transcript quantification, the biomarkers extracted from our *P. knowlesi* bulk RNA-seq time course can refine the scRNA-seq UMAP plot annotations.

**Figure 4.**
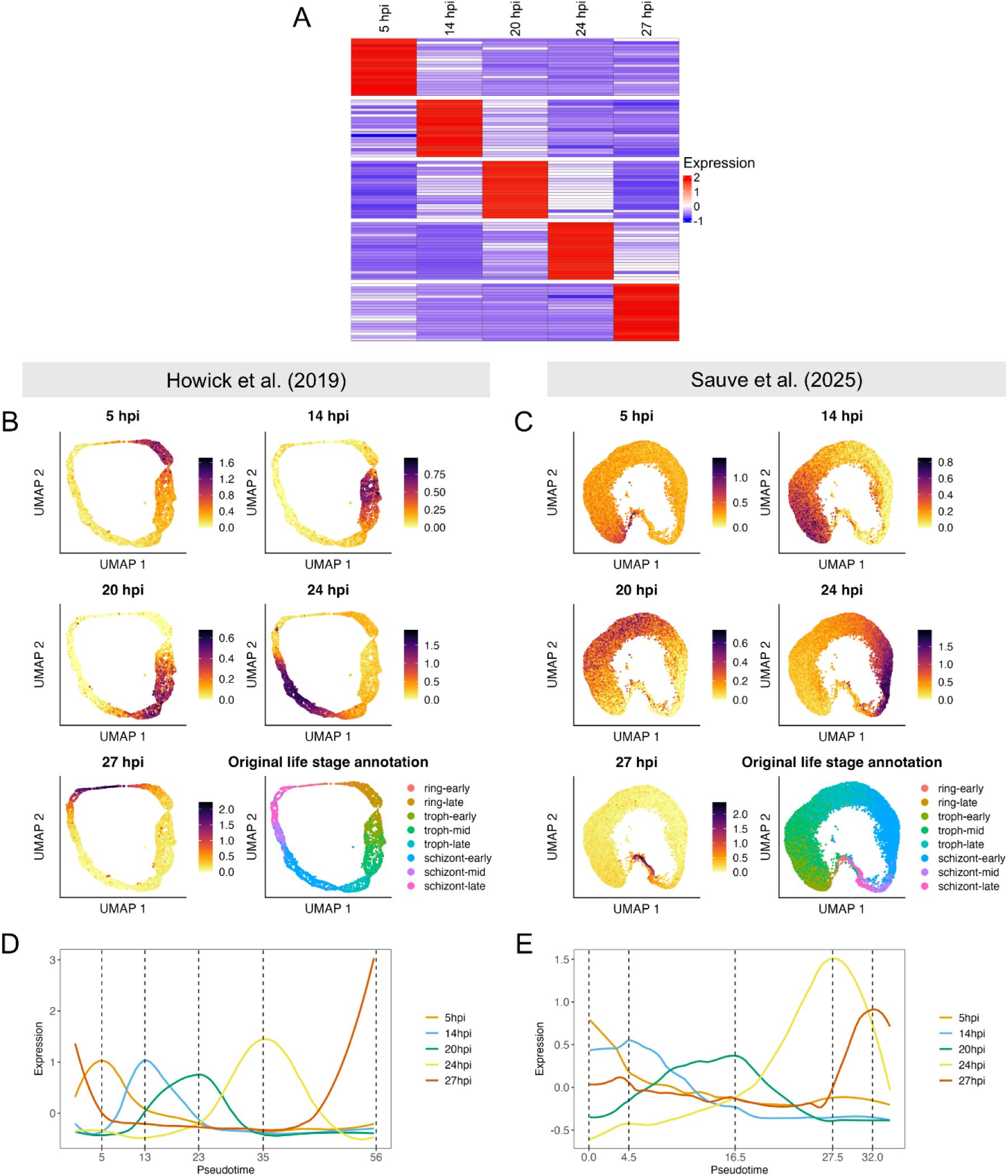
*P. knowlesi* stage-specific biomarkers and their use to annotate time points in single-cell datasets. **A)** Heatmap of stage-specific biomarkers derived from the *P. knowlesi* A1-H.1 transcriptome time course, showing the expression (TPM-normalised, Z-scored) of the 20 biomarker genes (rows) that were selected for each time point (columns). Per time point, genes with a TPM-normalised expression that was >3-fold upregulated compared to every other time point were identified, from which the 20 genes with the highest TPM-normalised expression at that time point were selected as biomarkers. **(B-C)** Time point annotation of two publicly available *P. knowlesi* A1-H.1 single-cell RNA-seq datasets, generated using a 10X Genomics (27) (B) and a HIVE (28) (C) platform. For each time point, the average single-cell gene expression of the 20 stage-specific biomarker genes was visualised on a UMAP. The ‘Original life stage annotation’ panel shows life cycle stage annotation based on a *P. berghei* microarray time course as done by Howick et al. (2019) (27), which was later transferred to the HIVE data from Sauve et al. (2024). **(D-E)** Expression of biomarkers over pseudotime, shown for single-cell data from Howick et al. (2019) (D) and Sauve et al. (2024) (E). For every set of stage-specific biomarker genes, the average expression (scDESEQ2-normalised, Z-scored) of the 20 genes was calculated at every pseudotime-point and shown as a coloured and LOESS-smoothed line. The peak position of each set of stage-specific biomarkers can be linked to a pseudotime (dashed vertical lines).

When the average expression pattern of the 20 biomarkers is visualised in the single-cell UMAP plots derived from 10X Genomics (27) and HIVE (28) datasets (**Fig. 4**), distinct regions can be identified for each of the five time points, which follow each other in a manner consistent with the expected cyclic progression of the IDC. Life cycle stages were originally annotated by Howick et al. (2019) based on a *P. berghei* microarray time course (**Fig. 4B** panel ‘Original life stage annotation’). This annotation was subsequently projected on the single-cell data from Sauve et al. (2024) (**Fig. 4C** panel ‘Original life stage annotation’). Our annotation of *P. knowlesi* bulk RNA-seq time points to these single-cell datasets largely aligns with the prior life cycle stage annotations based on *P. berghei*, except for cells identified as late schizonts (27 hpi) in our time course dataset, which were partly categorised as early rings in the *P. berghei*-based annotation. However, we can confidently exclude the presence of ring stages in our 27 hpi late schizont sample, as a Nycodenz purification was applied to enrich schizonts before collection, and microscopy confirmed the absence of rings (**Fig. 1**). Therefore, our use of bulk RNA-seq data from tightly synchronised *P. knowlesi* A1-H.1 parasites refines the time point annotation of published single-cell transcriptome datasets and is particularly useful for distinguishing late schizont from ring cells transcriptomes. Furthermore, pseudotime analysis of single-cell data is often used to order parasite cells along a continuous developmental trajectory, thereby assigning a relative temporal position to each parasite cell. Our stage-specific biomarkers with known peak expression times can be used to infer the biological age (hpi) that corresponds to each pseudotime age (**Fig. 4D-E**), effectively anchoring the pseudotime trajectory to developmental time. Taken together, the biomarker genes identified here can be confidently used to annotate and calibrate future *P. knowlesi* A1-H.1 or *P. vivax* single-cell transcriptome datasets.

### Expression patterns of *P. knowlesi* large multigene families

The variant antigens encoding the Schizont-Infected Cell Agglutination proteins (SICAvar) represent the largest multigene family in *P. knowlesi* (33), containing 233 members in strain H. The SICAvar proteins are located on the surface of *P. knowlesi*-infected RBCs and mediate cytoadherence, influence parasite virulence and have a role in immune evasion (33–38). The *SICAvar* multigene gene family is expressed predominantly at low levels in *P. knowlesi* A1-H.1, with 203 genes exhibiting maximal expression levels that fall below the first quartile (25%) of the genome-wide distribution of maximal expression levels (**Fig. 5A**). In contrast, a small subset (n=9) of *SICAvar* genes displays higher maximal expression levels, exceeding the median or 3^rd^ quartile (75%) of the genome-wide distribution. Among these, PKNH_0620500 shows the highest expression, with 6.7-fold higher expression levels than the second most highly expressed *SICAvar* gene (PKNH_1021300). PKNH_0620500 expression peaks at the schizont stage (24 hpi). Other moderately highly expressed *SICAvar* genes often peak at the schizont stage as well, although other peak timings are also observed (**Fig. 5B**).

**Figure 5.**
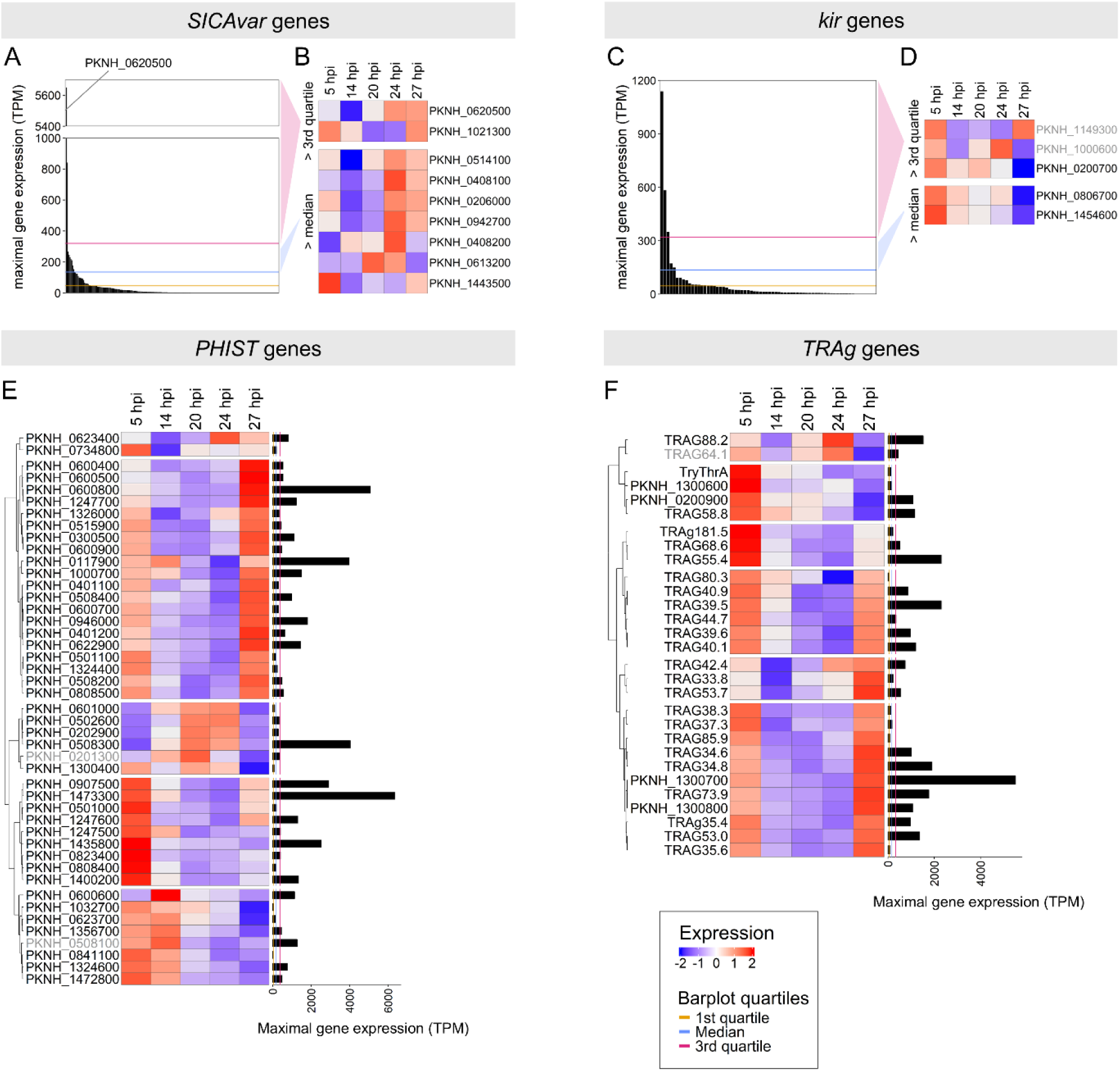
Expression of the *SICAvar*, *kir*, *PHIST* and *TRAg* multigene families. **(A, C)** Bar plots showing the maximal gene expression level (TPM-normalised) out of the 5 sampled time points for all members of the *SICAvar* or *kir* gene family. The y-axis of the *SICAvar* plot is interrupted to visualise the highly expressed gene PKNH_0620500. The yellow, blue and pink horizontal lines indicate the 1^st^ quartile, median and 3^rd^ quartile of the genome-wide distribution of maximal expression levels, respectively. **(B, D)** Heatmaps of normalised (TPM-normalised, log_2_-transformed and Z-scored) expression of the *SICAvar* or *kir* genes (rows) whose maximal expression exceeds the genome-wide median and 3^rd^ quartile, for the 5 collected time points (columns). Gene IDs displayed in grey indicate low transcriptional variation over the IDC (log_2_ fold change between highest and lowest expression value below the filtering threshold of 0.5). **(E-F)** Clustered heatmaps of all members of the *PHIST* or *TRAg* gene families, with bars reflecting their expression level on the side. Heatmaps show normalised (TPM-normalised, log_2_-transformed and Z-scored) expression of *PHIST* or *TRAg* genes (rows) for the 5 collected time points (columns). Gene IDs displayed in grey indicate low transcriptional variation over the IDC (log_2_ fold change between highest and lowest expression value (TPM, log_2_) below the filtering threshold of 0.5). Bars show the maximal gene expression level (TPM-normalised) out of the 5 sampled time points. The yellow, blue and pink vertical lines indicate the 1^st^ quartile, median and 3^rd^ quartile of the genome-wide distribution of maximal expression levels, respectively.

The *Plasmodium* interspersed repeat genes (*pir*) superfamily, also known in *P. knowlesi* as the *kir* family (39, 40), has been linked to immune evasion, pathogenesis and cytoadhesion in *P. vivax* and *P. knowlesi* (34, 41–43). In *P. knowlesi* A1-H.1, the majority of *kir* genes (53/70) show maximal expression levels below the first quartile of the genome-wide distribution of maximal expression levels. However, three *kir* genes (PKNH_1149300, PKNH_1000600, PKNH_0200700) show higher maximal expression above the genome-wide third quartile (**Fig. 5C**). Unlike the *SICAvar* genes, these three highly transcribed *kir* genes show comparable expression levels. Their peak expression occurs at different time points, although expression during the ring stage is common (**Fig. 5D**).

The last two large multigene families that were investigated are *Plasmodium* Helical Interspersed Sub-Telomeric (*PHIST)* and Tryptophan-rich Antigen (*TRAg*). The *PHIST* family exists across the *Plasmodium* genus, and PHIST proteins are exported to the cytoplasm, where they are implicated in RBC remodelling (44, 45). *PkTRAg* family genes have been implicated in RBC binding and invasion (46, 47). Unlike *SICAvar* and *kir*, the *PHIST* and *TRAg* families show a large proportion of highly expressed genes (32/45 genes and 20/29 >3^rd^ quartile genome-wide, respectively) (**Fig. 5E, F**). In both gene families, distinct temporal expression patterns over the IDC can be distinguished, although *TRAg* genes peak predominantly during the late schizont and/or ring stages (**Fig. 5E, F**).

### Comparison of temporal expression patterns between *P. vivax* – *P. knowlesi* orthologues

The close phylogenetic relationship of *P. vivax* to *P. knowlesi* allows comparison of transcriptional dynamics between *P. vivax* genes (publicly available *ex vivo* time course) (31) and their *P. knowlesi* orthologues over the IDC. As the two species have different IDC lengths (*P. knowlesi* 27h, *P. vivax* 48h), IDC time was first normalised to a 0-1 scale for both species. To quantify the similarity between orthologue gene expression patterns, we used a combination of a dynamic time warping-based similarity score and the difference in peak expression timing between the orthologues. Out of the 3877 orthologous gene pairs that were assessed, 2920 pairs (75%) were classified as having similar temporal expression patterns (‘similar orthologues’) (**Fig. 6A, B**), while the remaining 957 pairs (25%) were classified as dissimilar (‘dissimilar orthologues’) (**Fig. 6A, C**; classification in **Additional File 4**). Since the classification of orthologous gene pairs as similar or dissimilar depends on arbitrarily selected parameters (*e.g.* temporal shift allowed between the orthologues’ expression patterns), an interactive web tool was developed in which the temporal expression of any *P. vivax* – *P. knowlesi* orthologous gene pair can be visualised and assessed individually. In addition, the temporal expression pattern of any *P. knowlesi* or *P. vivax* gene of choice can be explored (https://interactive.itg.be/app/mal-pk-pv-expression-viewer).

**Figure 6.**
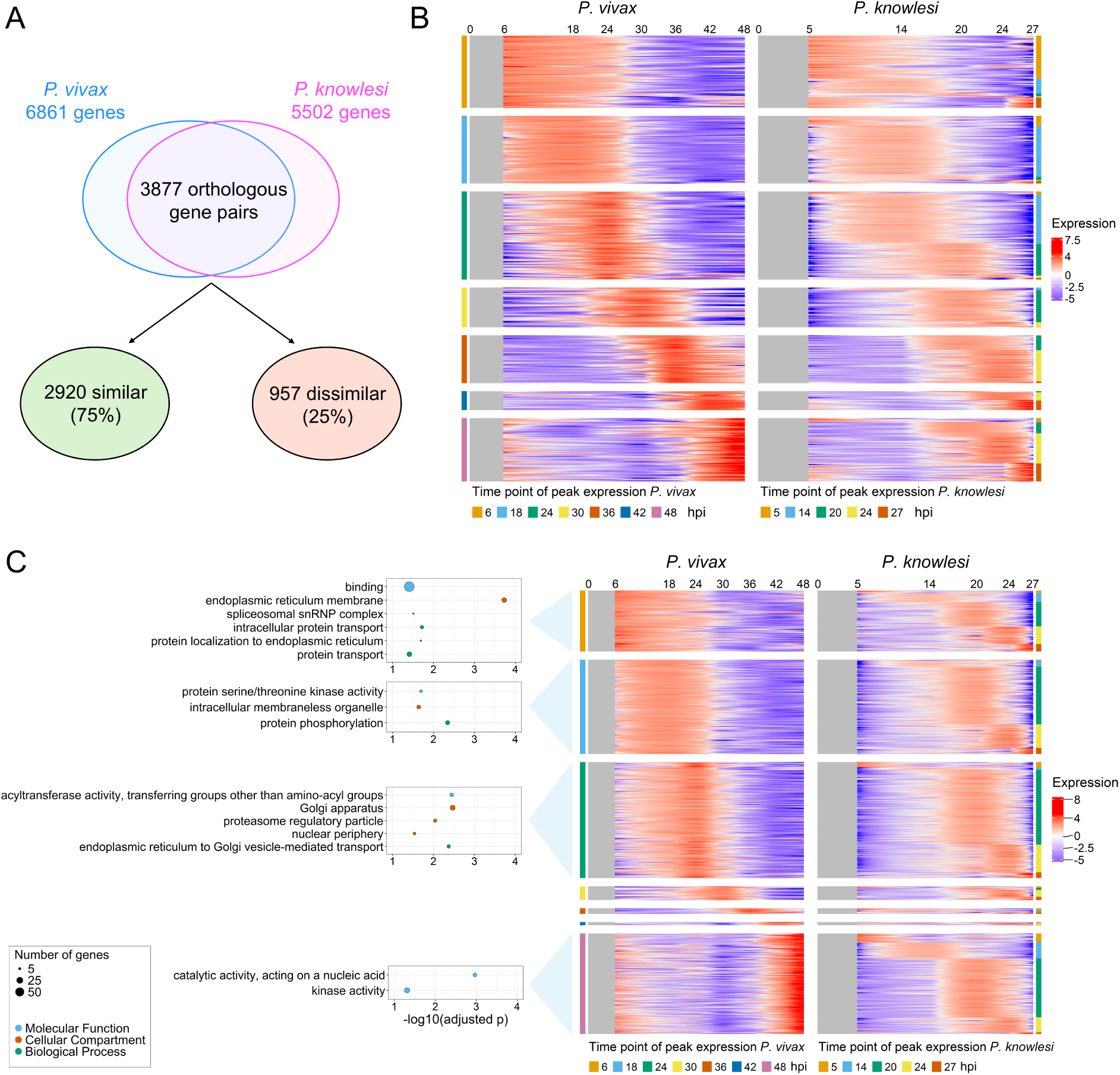
*P. vivax – P. knowlesi* orthologous gene pairs with ‘(dis)similar’ temporal expression patterns over the IDC. **(A)** Venn diagram showing the number of orthologous gene pairs classified as having similar or dissimilar temporal expression patterns. **(B)** Heatmap of normalised (TPM-normalised, log₂-transformed, Z-scored) and PCHIP-interpolated expression levels (rows) of *P. vivax* – *P. knowlesi* orthologue pairs with a temporal expression pattern classified as similar, for the full IDC (columns). Sampled IDC time (hpi) is indicated above the columns; intermediate time points are interpolated. *P. vivax* data originate from a published field isolate RNA-seq dataset (31). Only orthologue pairs for which both genes showed sufficient expression and temporal variability in the discrete data (filtering criteria detailed in Methods) were included. Grey shading indicates IDC intervals lacking data (no interpolation possible). Genes were firstly grouped by the peak expression time point of the *P. vivax* gene and, and within these time point blocks, ordered by the peak expression time point of the *P. knowlesi* gene. Peak expression time points for *P. vivax* and *P. knowlesi* genes are shown in coloured bars on the left and right, respectively. **(C)** Heatmap as in (B), now showing orthologue pairs with a temporal expression pattern classified as dissimilar. For each group of ‘dissimilar’ *P. vivax* genes (grouped by peak expression time point), gene ontology (GO) enrichment analysis was performed against a background of all *P. vivax* genes (classified as ‘similar’ and ‘dissimilar’) with a *P. knowlesi* orthologue peaking at the same time point. Bubble plots display significantly enriched GO terms per time point (Bonferroni-adjusted p<0.05). The x-axis shows the negative logarithm of the p-value, with higher numbers indicating higher significance. Bubble size reflects gene number; colours indicate GO categories (Molecular Function, Cellular Component, Biological Process). If no category or plot is shown, no significant enrichment was detected.

The majority of the orthologous gene pairs show a similar expression pattern in *P. vivax* and *P. knowlesi*, suggestive of a similar function and the possibility to extrapolate *P. knowlesi* experimental findings to *P. vivax*. For 25% of the orthologous gene pairs, *P. knowlesi* may not be an appropriate *P. vivax* model because expression patterns, and thus functions, are probably different. The function of these ‘dissimilar orthologues’ could reveal species-specific biological differences. To explore this, a GO enrichment analysis was performed on the ‘dissimilar’ *P. vivax* genes, comparing them per peak expression time point to all *P. vivax* genes (both ‘similar’ and ‘dissimilar’ with a *P. knowlesi* ortholog) peaking at that same time point (**Fig. 6C**). This revealed that at 6 hpi, *P. vivax* ring-stage ‘dissimilar’ genes are enriched for functions related to the endoplasmic reticulum, protein trafficking and RNA splicing. At 18 hpi and 48hpi, ‘dissimilar’ *P. vivax* genes are predominantly linked to catalytic activities. Finally, ‘dissimilar’ *P. vivax* genes peaking at 24 hpi display a diverse set of functions, including protein secretion and degradation.

### Comparing expression patterns of invasion-associated orthologues between *P. knowlesi* and *P. vivax*

To assess the validity of *P. knowlesi* as a model to study the *P. vivax* RBC invasion process, we compared the temporal expression dynamics of invasion-related genes between the two species. We focused on invasion-related genes that have orthologues between *P. knowlesi* and *P. vivax* (regardless of their classification as ‘similar’ or ‘dissimilar’). The *DBP* genes were included because PvDBP and PkDBPα are ligands of the human Duffy Antigen Receptor for Chemokines (DARC), an essential interaction for invasion in both species (48–50), while PkDBPβ and PkDBPγ bind to unidentified macaque RBC receptors (51, 52). The *P. vivax* Reticulocyte Binding Proteins (RBP) family consists of 10 members (genome PvP01) and is orthologous to the two-member *P. knowlesi* Normocyte Binding Protein X (NBPX) family. *Pv*RBP2a and *Pv*RBP2b are ligands of reticulocyte receptors CD98 and CD71, respectively (53, 54), while *Pv*RBP1a interacts with CD71, prohibitin-2 and basigin (55). The *TRAg* multigene family is encoded by 29 members in the *P. knowlesi* genome (strain H) and 40 members in the *P. vivax* genome (PvP01), sharing 26 orthologues. Multiple TRAg’s from both *P. knowlesi* and *P. vivax* show erythrocyte binding activity or have a role in invasion (46, 47, 56–58). In addition, *Rhoptry Neck Protein (RON)* genes (59–61), *MSP* genes (62–65), *GPI-anchored Micronemal Antigen (GAMA)* (66, 67), *merozoite-specific thrombospondin-related anonymous protein (MTRAP)* (68), *Apical Membrane Protein 1 (AMA1)* (61, 69), *Cysteine-rich Protective Antigen (CyRPA)* (70) and *Rh5 Interacting Protein (RIPR)* (70) were included.

The temporal expression patterns of these invasion-associated genes are largely conserved between *P. knowlesi* and *P. vivax*, confirmed by the majority of orthologues being classified as similar (**Fig. 7A**). As expected, expression generally peaks during the schizont stage (genes in clusters 2-6, **Fig. 7A**). Clusters 2-3 (including mainly *MSP* and *RON* genes) show earlier schizont stage expression, while genes in clusters 4-6 (including *TRAg* and *DBP* genes among others) as well as the orthologous families *PkNBPX/PvRBP* genes show late schizont expression with additional ring stage expression (cluster 4, *P. knowlesi* cluster 5 and 6). In contrast, cluster 1 (which also includes some genes from the *TRAg* family and *MSP8*) shows predominant ring stage expression, implying that these genes are unlikely to participate in invasion. Despite the overall conserved expression patterns between both species over the IDC, differences are also observed. For example, ring stage expression in genes in cluster 5 and 6 is observed in *P. knowlesi* but not in *P. vivax*; and schizont stage expression begins slightly earlier in *P. vivax* for genes in clusters 4 and 5, which include genes such as *DBP*s, *AMA1*, *RIPR*, *CyRPA* and *GAMA*, and the same can be observed for the *PkNBPX/PvRBP* family.

**Figure 7.**
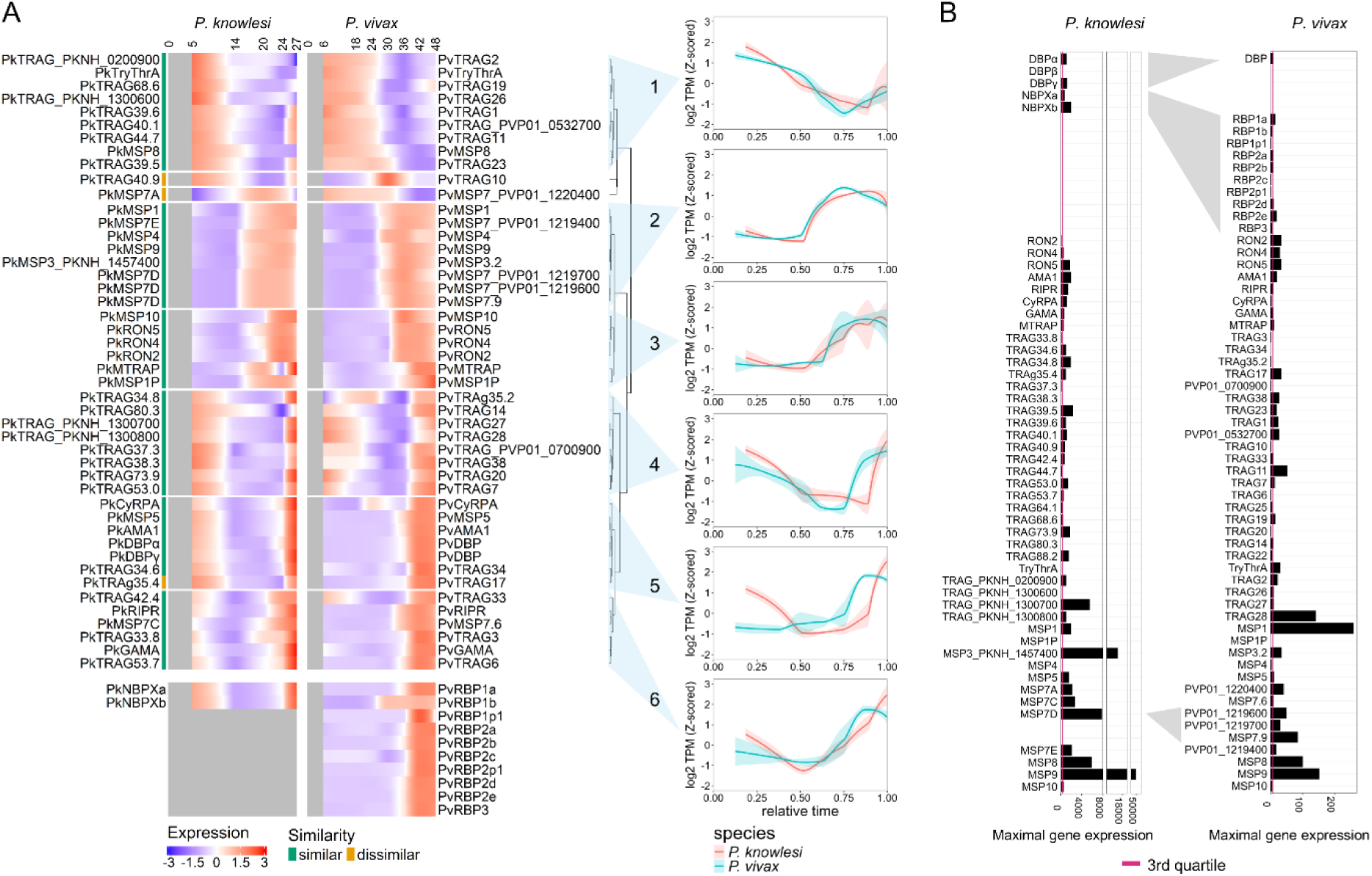
Comparison of expression patterns of invasion-associated *P. knowlesi* – *P. vivax* orthologous gene pairs. *P. vivax* data originate from a published field isolate RNA-seq dataset (31). Invasion-associated genes with no orthologue in *P. knowlesi* or *P. vivax* are not shown. **(A)** Heatmap of normalised (TPM-normalised, log_2_-transformed, Z-scored) and PCHIP-interpolated expression levels (rows) of *P. vivax* – *P. knowlesi* orthologue pairs associated with invasion, for the full IDC (columns). Sampled IDC time (hpi) is indicated above the columns; intermediate time points are interpolated. Orthologue pairs for which one of the members did not pass the filtering (too low expression or temporal variability in the discrete data, see Methods) were excluded (*DBPβ*, *PkTRAg64.1*, *PvTRAg25*, *PvTRAg22*). *PvDBP* and *PkMSP7D* appear multiple times, since they are part of multiple orthologue pairs. The coloured bar on the left indicates whether each orthologue pair was classified as similar or dissimilar in expression pattern. Grey shading indicates IDC intervals lacking data (no interpolation possible). Clustering was applied on both the *P. knowlesi* and *P. vivax* expression patterns together. For each of the 6 resulting clusters, a line plot is provided showing the mean *P. knowlesi* and *P. vivax* expression pattern (TPM-normalised, log-transformed, Z-scored) for that cluster with 95% confidence interval. Because the *PkNBPX* and *PvRBP* orthologous gene families lack one-to-one orthologous gene pairs, their members are shown separately at the bottom of the heatmap. **(B)** Bar plots showing the maximal gene expression level (TPM-normalised) out of the sampled time points for all invasion-associated genes (no gene filtering). Genes are ordered per family, and orthologous genes align horizontally (if a gene is orthologous to multiple genes, this connection is indicated by a grey field). The pink vertical line indicates the 3rd quartile of the genome-wide distribution of maximal expression levels, calculated separately for *P. knowlesi* and *P. vivax*.

Additionally, most invasion-associated *P. knowlesi* genes are highly expressed, and their maximal expression levels lie well above the 3^rd^ quartile (75%) of the genome-wide distribution of maximal expression levels (**Fig. 7B**). The human DARC ligand *DBPα* is highly expressed (71, 72), but so is *DBPγ*, even though it is a ligand for macaque RBCs (51, 52) and parasites were grown exclusively in human RBCs. Transcripts for *DBPβ*, another macaque RBC ligand (51, 52), were not detected due to a gene deletion in the A1-H.1 clone used in this study (**Supplementary Fig. S1A-B**). Similarly, both *NBPXa* and *NBPXb* are highly expressed, and although only *NBPXa* can bind to human RBCs within the *PkNBPX* family (73, 74), *NBPXb* is expressed at higher levels. *P. vivax* invasion-associated genes generally show lower expression levels than *P. knowlesi* when each gene is compared to its own 3^rd^ quartile. Orthologues of the *TRAg* family show variation in expression levels between both species, and *MSP* genes range from the very low expression levels of *MSP1P* up to the highest expression levels of invasion-associated genes (*PkMSP9*, *PvMSP1*).

### Comparing ApiAP2 transcription factor expression patterns between *P. knowlesi*, *P. vivax* and *P. falciparum*

ApiAP2 transcription factors are key regulators of progression through the IDC, and have been studied extensively in *P. falciparum* (23–26). In *P. knowlesi*, 30 ApiAP2 transcription factor-encoding genes are annotated, which display stage-specific expression as in other *Plasmodium* species (**Fig. 8A**).

**Figure 8.**
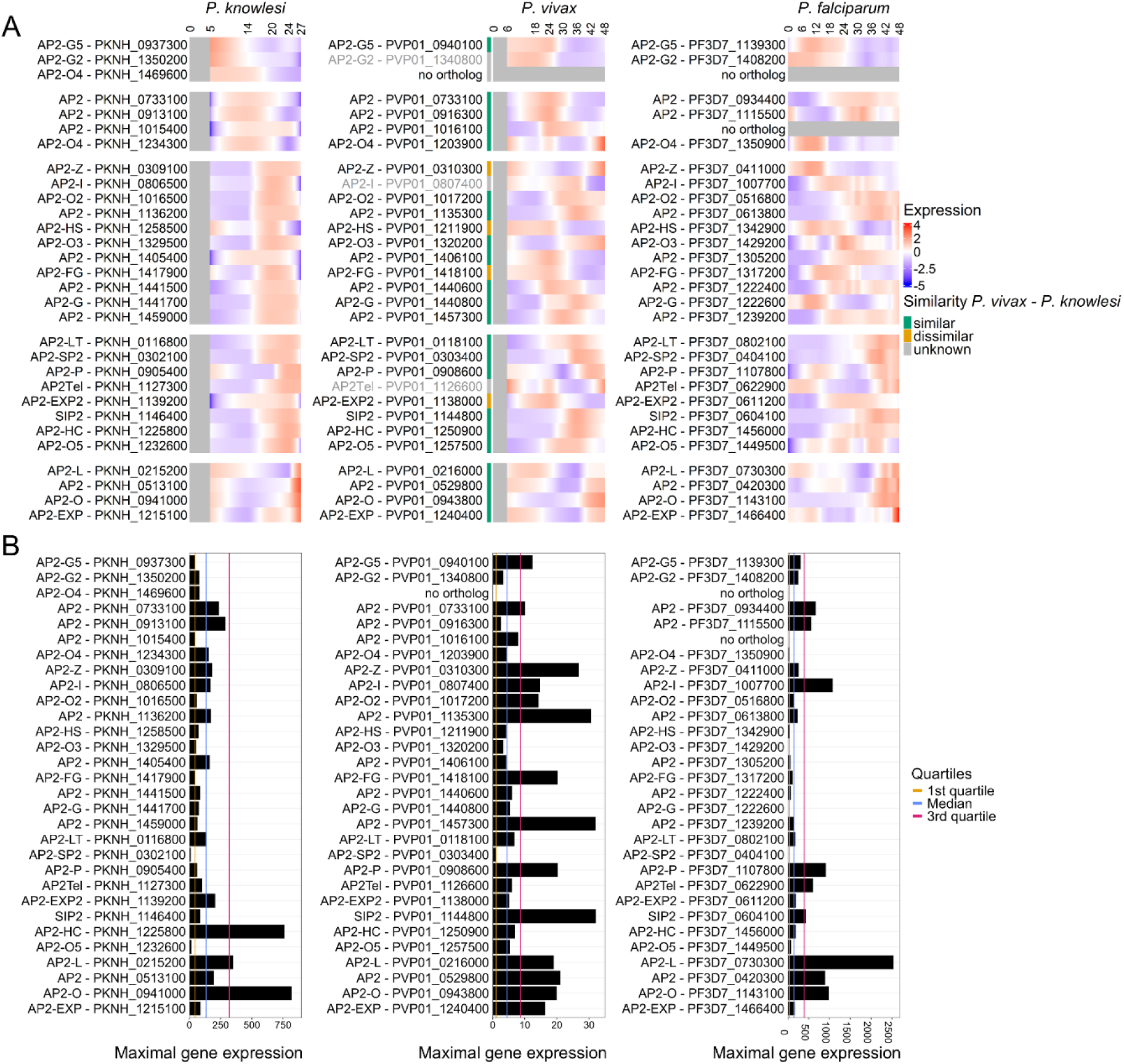
Comparison of ApiAP2 transcription factor expression patterns between *P. knowlesi*, *P. vivax* and *P. falciparum*. *P. vivax* and *P. falciparum* data originate from published RNA-seq datasets (31, 90), derived from a field isolate and laboratory strain (II3, 3D7-derived), respectively. **(A)** Heatmap of normalised (TPM-normalised, log_2_-transformed, Z-scored) LOWESS-smoothed (only *P. falciparum*) and PCHIP-interpolated expression levels of all *P. knowlesi ApiAP2* genes and their orthologues in *P. vivax* and *P. falciparum* (rows, orthologues are horizontally aligned), for the full IDC (columns). For *P. knowlesi* and *P.* vivax, sampled IDC time (hpi) is indicated above the columns; intermediate time points are interpolated. The *P. falciparum* time course was sampled every 2 hours, interpolated for intermediate time points, and labelled every 6 hours for readability. Genes are ordered by their peak expression timing in *P. knowlesi*, with horizontal gaps splitting *P. knowlesi ApiAP2* genes peaking at 5 hpi, 14 hpi, 20 hpi, 24 hpi or 27 hpi. Genes that did not pass the filtering (too low transcriptional variability in temporal pattern in the discrete data, see Methods) are shown as grey text. The coloured bar left of the *P. vivax* heatmap indicates whether each *P. knowlesi* – *P. vivax* orthologue pair was classified as similar or dissimilar in expression pattern. Grey shading indicates IDC intervals lacking data (no interpolation possible), or absence of an orthologue. **(B)** Bar plots showing the maximal gene expression level (TPM-normalised) out of the sampled time points for all *P. knowlesi ApiAP2* genes and their orthologues in *P. vivax* and *P. falciparum*. Genes are in the same order as in which they appear in the heatmaps (A), and orthologous genes align horizontally. The yellow, blue and pink vertical lines indicate the 1^st^ quartile, median and 3^rd^ quartile of the genome-wide distribution of maximal expression levels, respectively, calculated separately for the 3 species.

Although the majority of *P. knowlesi ApiAP2* genes exhibited temporal profiles consistent with those of their *P. vivax* and *P. falciparum* orthologues, differing expression patterns were also observed (**Fig. 8A**). *PkAP2-HS* (regulator of the protective heat-shock response in *P. falciparum*) (75), *PkAP2-Z* (critical for zygote development in *P. berghei*) (76) and *PkAP2-FG* (female gametocyte development in *P. berghei*) (77) differed from both other species; *PkAP2-EXP2* (involved in RBC remodelling in *P. falciparum*) (78) and *PkAP2-O4* (ookinete development) (26) differed from *P. vivax*; and *PkAP2-I* (involved in invasion in *P. falciparum*) (79), *PkAP2Tel* (co-localises with telomeric clusters at the nuclear periphery in *P. falciparum*) (80), *PkAP2-O3* (regulator of female gametocytes in *P. yoelii* and *P. berghei*) (81, 82), *PkAP2-O5* (ookinete development) (26), *PkAP2-G* (master regulator of sexual conversion) (83–85), *PKNH_0733100* and *PKNH_0913100* differed from *P. falciparum* (transcriptional variability in temporal pattern of *PvAP2-I* and *PvAP2Tel* was too low to compare to *P. knowlesi*) (**Fig. 8A**).

When comparing maximal expression levels, *P. knowlesi ApiAP2* transcription factors are generally expressed at lower levels than their *P. vivax* and *P. falciparum* orthologues. Only 43% of *P. knowlesi ApiAP2* transcription factors have maximal expression above the genome-wide median, compared with 76% in *P. vivax* and 68% in *P. falciparum* (**Fig. 8B**). However, *PkAP2-HC* (associates with heterochromatin in *P. falciparum*) (26) is very highly expressed compared to both other species. Some *PkApiAP2* genes show very low expression levels (*AP2-SP2*, *AP2-O5*), consistent with their role in non-erythrocytic life stages (26), although, other *PkApiAP2* genes linked to non-erythrocytic stages remain relatively highly expressed during the IDC (*e.g. AP2-L*, *AP2-O*, *AP2-O4*, *AP2-Z*), suggesting potentially additional functions during the asexual blood stage development in *P. knowlesi* (26).

Although *P. knowlesi* line A1-H.1 does not produce gametocytes (11), we assessed the expression pattern of *PkAP2-G*, the conserved master regulator of sexual conversion in *P. falciparum*, *P. berghei* and *P. yoelii* (84, 86, 87). *PkAP2-G* revealed peak expression in the late trophozoite to mid-schizont stage (20-24 hpi) and shows no detectable ring stage expression, whereas *P. falciparum AP2-G* expression peaks in late schizonts and the ring stage (**Fig. 8A**) (88, 89). Interestingly, the *PkAP2-G* temporal expression pattern resembles that of *P. vivax AP2-G* better (**Fig. 8A**): neither species show detectable ring stage *AP2-G* transcripts, while in *P. falciparum PfAP2-G* is first detected in sexual rings (the stage of sexual development that follows the committed schizont stage) (88). *AP2-G* reaches a higher maximal expression in *P. knowlesi* than in *P. falciparum*, though still has a lower maximal expression than in *P. vivax*, when compared to each species’ genome-wide distribution of maximal gene expression (**Fig. 8B**).

Intriguingly, only the 3’ end of *PkAP2-G* appears to be transcribed; the 5’ end shows no detectable signal (**Supplementary Fig. S2A**). This is different from *P. vivax*, where full-length *PvAP2-G* expression was observed (**Supplementary Fig. S2D**). We observed no *PkAP2-G* deletion or high-confidence mutations in the promoter or gene (**Supplementary Fig. S2B**), absence of transcripts covering the 5’ end was not observed in other genes (including large genes with a similar size to *PkAP2-G*; **Supplementary Fig. S2C**), and the *AP2-G* regulator *GDV1* is fully expressed without mutations in its coding region or promoter (**Supplementary Fig. S3A-B**). Unexpectedly, an antisense transcript covering the 3’ UTR was observed in *P. knowlesi* AP2-G, which has not yet been reported for *P. falciparum* (**Supplementary Fig. S2A**).

## Discussion

In this study, we present the first transcriptome time course for the IDC of *P. knowlesi* line A1- H.1. Previously, a *P. knowlesi* transcriptome time course was constructed for clone Pk1(A+) in rhesus blood (21), comparing *in vitro* and *ex vivo* transcript levels and identifying multiple differentially expressed genes highlighting the impact of host context. In contrast, our time course focuses on the human-blood adapted line Pk A1-H.1 to increase comparability with *P. vivax*. From these data, we identified stage-specific biomarkers and constitutively expressed genes and systematically compared *P. knowlesi* expression patterns with their *P. vivax* orthologues. To facilitate broader accessibility and enable comparative analyses, we developed an interactive web tool that allows researchers to compare the expression profiles between orthologous genes in the two species, as well as any other gene of choice. Because expression pattern similarity serves as a proxy for likely functional conservation between both species, this tool enables rapid identification of which *P. knowlesi* orthologues are suitable to study gene function in *P. vivax*. Three quarters of the *P. vivax* genes with an orthologue in *P. knowlesi* (2920/3877) were classified as having expression profiles similar to their *P. knowlesi* orthologues (91). Given the phylogenetic proximity of both species, this level of concordance is expected and supports the suitability of Pk A1-H.1 as a model system to investigate *P. vivax* biology.

Our *P. knowlesi* A1-H.1 time course places a special emphasis on late stage time points (20-27 hpi), which are of particular interest due to the expression of invasion ligands. Those are crucial in life cycle progression and are attractive targets for the development of vaccines and therapeutic interventions (92–94). The invasion process of *P. knowlesi,* similar to *P. vivax,* remains elusive despite recent progress in elucidating the sequential steps of human RBC entry (95). Although invasion pathways are known to differ between *P. vivax*, *P. falciparum* and *P. knowlesi* (2, 96), largely due to different RBC tropisms and the high selective pressure on invasion ligands (97, 98), *P. knowlesi* still shares multiple invasion-associated genes with *P. vivax* (such as the *DBP, NBPX* and *TRAg* families). We show that almost all investigated invasion-associated genes share a similar temporal expression pattern between the two species. Only *PkMSP7A/PvMSP7* (*PVP01_1220400*), *PkTRAg35.4/PvTRAg17* and *PkTRAg40.9/PvTRAg10* showed expression patterns classified as ‘dissimilar’, and thus, likely have a divergent function between *P. knowlesi* and *P. vivax* or may reflect uncertainties in orthologue assignment within these multigene families. Furthermore, not all *Pv/PkTRAg* genes were schizont-expressed (20), indicating that despite their conserved Tryptophan/Threonine-rich domain, those *TRAg* family members likely have other functional roles than facilitating RBC invasion. It was previously hypothesised that ring-expressed *PvTRAg*’s may have a role in rosetting based on the RBC-binding capacity of certain *TRAg* proteins (57). Interestingly, a subgroup of *Pv*/*PkTRAg* genes was expressed in both the schizont and ring stage, which is suggestive of a dual function.

Overall, the temporal expression patterns of invasion genes are largely conserved between *P. knowlesi* and *P. vivax*. The minor differences observed likely reflect the inherent technical difficulties of comparing datasets from different species and different studies, rather than actual biological differences. For example, some invasion-associated genes (*e.g. DBP*, *NBPX/RBP* family) were also expressed at the ring stage in *P. knowlesi* A1-H.1, whereas they were not in *P. vivax.* This could be attributed to the presence of a small number of dead schizonts in our *P. knowlesi* 5 hpi sample. Furthermore, some *P. knowlesi* invasion-associated genes were transcribed later in the schizont stage compared to their *P. vivax* orthologues (*e.g. DBP* family, *PkNBPX/PvRBP* family, *TRAg* family members, *AMA1*, *CyRPA*, *RIPR*). This shift may be due to the lower synchrony of the *P. vivax* time course culture, inherent to the use of field isolates, suggesting that the true timing of *P. vivax* gene expression may be closer to that of *P. knowlesi* A1-H.1. Moreover, as *P. vivax* cannot be continuously cultured *in vitro*, data from an *ex vivo* clinical isolate may not be fully comparable to those from a culture-adapted *P. knowlesi* line. Finally, rescaling of the different IDC lengths of *P. knowlesi* and *P. vivax* to a 0-1 interval might not perfectly reflect the biological alignment of life cycle stages.

Beyond transcriptional timing, the expression levels of invasion genes can also provide valuable insights. The transcript levels of *P. knowlesi* invasion genes indicate which ligands are actively expressed in the human-adapted Pk A1-H.1 line. Unlike *P. vivax*, *P. knowlesi* infects both humans and macaques. This dual host tropism is reflected in the *DBP* and *NBPX* invasion gene families: *DBPα* is a ligand for the human DARC receptor (71, 72), *DBPβ* and *DBPγ* interact with (an) unknown macaque RBC receptor(s) (51, 52), *NBPXa* binds unidentified human and macaque receptors (73, 74, 99) and *NBPXb* binds an unidentified macaque receptor (74, 99). Although Pk A1-H.1 was adapted to grow in human RBCs, the original line reportedly retained the capacity to replicate in macaque RBCs (11). Consistent with this, we observe *DBPγ* expression levels comparable to those of *DBPα*, and *NBPXb* expression even exceeding *NBPXa*. In contrast, *DBPβ* is transcriptionally silent, consistent with A1-H.1 single-cell RNA-seq data from Sauve et al. (2024), due to a genomic deletion spanning *DBPβ* and an upstream pseudogene. This deletion was not yet present in the original line Pk A1-H.1 (11, 100), suggesting that it emerged over the course of continuous culturing in human RBCs and might prevent the ability to grow in macaque RBCs. Furthermore, a comparison between invasion ligand expression in our human-adapted Pk A1-H.1 line and the Pk1(A+) clone grown in rhesus blood (21) would be highly informative to assess the impact of host species blood on *P. knowlesi* transcriptional regulation. However, such a comparison is complicated by the use of different sequencing platforms (RNA-seq versus microarray), making it difficult to disentangle technical from biological variation.

In addition to invasion genes, we also examined the expression of *PkApiAP2* transcription factors. The *PkApiAP2* genes displayed stage-specific expression that generally corresponded to those of their *P. vivax* or *P. falciparum* orthologues, although a substantial subset (11/29) showed divergent patterns that may warrant further investigation. Interestingly, even though *P. knowlesi* A1-H.1 does not produce gametocytes *in vitro*, it showed schizont-stage expression of *AP2-G*, the master regulator of sexual conversion in *P. falciparum*, *P. berghei* and *P. yoelii* (84, 86, 87). However, this transcription was restricted to the 3’ end of the coding sequence, suggestive of an alternative internal start site. The absence of full-length transcription may prevent production of a complete, functional PkAP2-G protein, and hence sexual conversion. This is consistent with the lack of *PkAP2-G* ring stage expression, which in *P. falciparum* follows schizont commitment and is a marker of sexual rings (88). On the other hand, *P. vivax AP2-G* does not show detectable ring stage expression either, even though *PvAP2-G* is fully expressed and gametocytes are expected in the clinical *P. vivax* isolate analysed. Whether sexual conversion and sexual differentiation in *P. knowlesi*, and possibly *P. vivax*, proceeds through the activation of *AP2-G* as master regulator of sexual conversion or via a distinct mechanism is not yet known and remains to be investigated.

Finally, expression patterns of large *P. knowlesi* multigene families were characterised over the IDC. The *P. knowlesi SICAvar* gene family is of particular interest as it plays an important, though often still elusive, role in *Plasmodium* host-parasite interactions (101). While the *SICAvar* family has no counterpart in *P. vivax*, it shares an evolutionary origin with the *P. falciparum var* gene family (35, 102). In our A1-H.1 dataset, the majority of the *SICAvar* genes were expressed at low levels, consistent with previous observations *in vitro* in the Pk1(A+) time course in rhesus RBCs of Lapp et al. (2015) and in splenectomised rhesus monkeys (37), but in contrast to *ex vivo P. knowlesi* infections in non-splenectomised rhesus monkeys (21). Similar to the *var* family, *SICAvar* genes show antigenic variation and are involved in cytoadherence (34, 38), which is not required under *in vitro* culture conditions, likely explaining their generally low expression in our *in vitro* time course. Nevertheless, a subset of *SICAvar* genes (notably PKNH_0620500) showed moderate to high expression, suggestive of mutually exclusive expression, or of divergent functions for some highly expressed *SICAvar* genes. Similarly, the *kir* genes show overall low expression levels, with a few highly expressed genes at the ring or schizont stage, indicative of clonally variant gene expression. The *P. vivax vir* genes, which are also part of the *pir* superfamily, indeed show transcriptional variability between field isolates at both the schizont (58, 103) and ring stage (31, 104). Consistent with this, HP1-enriched chromatin regions, markers of heterochromatin and associated with clonally variant gene expression, have been identified in *SICAvar* and *kir/vir* genes (105). The expression timing of highly expressed *kir* genes at either the schizont or ring stage is comparable to the *vir* genes, which undergo two ‘waves’ of antigenic presentation at these same time points (20). Nevertheless, conclusions regarding the temporal expression patterns and expression levels of *P. knowlesi* multigene families involved in invasion, cytoadherence, immune evasion or other processes subject to environmental selective pressure, should be validated in (synchronous) natural human or macaque infections or experimentally in non-splenectomised rhesus monkeys. The suspected clonally variant gene expression patterns should be confirmed at the single-parasite level.

## Conclusions

The *P. knowlesi* A1-H.1 transcriptome time course presented here provides a resource to advance our understanding of *P. knowlesi* human infections, and serves as a tool to identify *P. vivax* orthologous genes with similar expression patterns that can be studied in *P. knowlesi* line A1-H.1. A novel bioinformatic pipeline was used to characterise this conservation of temporal expression patterns, which can be explored through an interactive web tool. Together with the ability to generate transgenic *P. knowlesi* parasites, this tool can enhance the *in vitro* testing of vaccines and therapeutics for *P. knowlesi* and *P. vivax*. In addition, constitutively expressed genes and stage-specific biomarkers were identified from this dataset. While this study focused predominantly on late asexual stage time points, future time courses could increase resolution during the ring and trophozoite stages to better capture earlier asexual parasite stage transcription, or explore sexual development by using a gametocyte-producing *P. knowlesi* macaque line. To strengthen direct cross-species comparisons of expression dynamics between *P. vivax* and *P. knowlesi*, a more synchronous *P. vivax* time course would be valuable. Together, the Pk A1-H.1 transcriptome time course data and comparative web tool provide a resource for comparative functional studies across malaria parasites and support translational research in *P. vivax* and *P. knowlesi*.

## Methods

### Parasite culturing

*P. knowlesi* line A1-H.1 was grown at parasitaemia’s ranging from 0.2%-6%, and was maintained at 3% haematocrit in culture medium consisting of RPMI-1640 (Westburg Life Sciences) supplemented with 0.32 g/L sodium bicarbonate, 5.0 g/L Albumax II (Invitrogen, cat: 11021-029), 2.0 g/L D-glucose, 25 mM HEPES, 0.05g/L hypoxanthine, 0.005 g/L gentamicin, and 2mM L-glutamine with 10% (v/v) heat-inactivated horse serum (Invitrogen), as described previously (11). Parasite cultures were incubated at 37°C on a shaking plate, under atmospheric conditions of 90% N_2_, 5% O_2_ and 5% CO_2_.

### Parasite preparation and collection of parasite samples at 5 time points across the IDC

Parasites were first synchronised to a 1-hour time window: schizonts and late trophozoites were purified using 55% isotonic Nycodenz bed, washed and allowed to reinvade for 1 hour at 37°C shaking, after which a second 55% isotonic Nycodenz schizont purification was performed. This time the RBC pellet containing ring-infected and uninfected RBCs was selected. This time point, after the 1-hour reinvasion time, is designated as 1 hpi and the cultures were allowed to grow until harvest time (5, 14, 20, 24 and 27 hpi) (**Fig. 1**). The same synchronisation procedure was performed independently to obtain parasites at i) 5 hpi, ii) 14 hpi, or iii) 20, 24 and 27 hpi.

Although the obtained 1 hpi rings were tightly synchronised to a 1-hour time window, a few schizonts were still present due to carryover during the second Nycodenz purification step. To block merozoite invasion from those schizonts, polysodium 4-styrenesulfonate (PSS; 200µg/mL final concentration) was added to the culture flasks that were matured to 14, 20, 24 and 27 hpi. For the flask that would be matured to 5 hpi, schizonts that remained after reinvasion were eliminated using guanidine hydrochloride (GuHCl) at 1 hpi, to prevent that some would still be present as mature schizonts at the 5 hpi harvest time point. The 1 hpi culture was mixed with 20 pellet volumes of buffered 140mM GuHCl for 20 minutes at 37°C as described (106). Next, the culture was washed twice in RPMI (650g, 5min), after which the RBC pellet top fraction containing dead schizonts was removed. The bottom fraction, containing rings and uninfected RBCs, was placed back in culture for 4 additional hours until the 5 hpi harvest point.

Parasites or RBC pellets harvested at their specific hpi (**Table 1**), were preserved in RNAprotect (Qiagen) according to the manufacturer’s instructions, and stored at −80°C. The 20, 24 and 27 hpi cultures with late stages were loaded on a 55% isotonic Nycodenz bed to purify the late trophozoite or schizont fraction for harvest, and to eliminate young rings from the 27 hpi sample. The 5 hpi and 14 hpi ring and trophozoite cultures were stored in RNAprotect (infected+uninfected RBCs) without prior parasite enrichment.

**Table 1.** Properties of samples used for RNA extraction and their resulting RNA concentration. Enrichment was done using a 55% isotonic Nycodenz schizont purification. Parasitaemia was assessed by light microscopy. RNA concentration was measured with the Qubit HS RNA kit.

| | Parasite enrichment | Parasitaemia | RBC pellet volume (μL) | RNA concentration (ng/μL) |
| --- | --- | --- | --- | --- |
| **5 hpi** | NA | 3.1% | 350 | 22.8 |
| **14 hpi** | NA | 2.9% | 550 | 54.2 |
| **20 hpi** | Nycodenz | ~90% | 9 | 102.0 |
| **24 hpi** | Nycodenz | ~90% | 9 | 53.5 |
| **27 hpi** | Nycodenz | ~90% | 3.5 | 43.1 |

RNA was extracted from RNAprotect-stored samples using the RNeasy Mini Kit (Qiagen, #74104) following the manufacturer’s instructions. In brief, samples were pelleted at high speed (20 min, 14,000 g), the supernatant was removed, and the pelleted cells were disrupted by adding 600 µL (20, 24, 27 hpi samples) or 1200 µL (5, 14 hpi samples) of RLT buffer. An on-column DNA digestion was performed using the RNAse-Free DNase Set (Qiagen, #79254). Elution in 30 µL of water resulted in RNA concentrations of 23-102 ng/µL (**Table 1**).

### mRNA-sequencing

RNA-extracted samples (20µL, containing 0.5-2.0 µg of RNA) were sent to GenomeScan B.V. (Leiden, The Netherlands) for library preparation and bulk mRNA-sequencing. Library preparation was carried out with the NEBNext Ultra II Directional RNA Library Prep Kit for Illumina (New England Biolabs, #E7760S/L), and included mRNA enrichment with oligo-dT magnetic beads and depletion of human globin mRNA using the QIAseq FastSelect −Globin Kit (Qiagen, #334376). The 150 base pair paired-end Illumina sequencing on a NovaSeq6000 resulted in 11-21 million read pairs per sample (**Supplementary Fig. S4**). The obtained fastq files are publicly available in the European Nucleotide Archive (ENA; https://www.ebi.ac.uk/ena/) under Study PRJEB83831.

### Read mapping and transcriptome analysis

For each sample, mRNA reads were aligned to the *P. knowlesi* strain H reference genome (PlasmoDB v2 release 66) and number of reads per gene were counted using STAR v2.7.6 (107). 88%-95% of all read pairs per sample mapped to the *P. knowlesi* genome, with the majority mapping to unique genomic locations (**Supplementary Fig. S4**). Human contamination was minimal (0.1%-4.2%; **Supplementary Fig. S4**). Aligned read coverage can be visualised in Integrated Genome Viewer (IGV) using the IGV session file available on Zenodo (https://doi.org/10.5281/zenodo.18608274). Coverage tracks displayed in IGV were created with deepTools (v3.5.5), and only primary alignments with a STAR mapping quality of 255 were included.

Next, a transcripts Per Million (TPM) normalisation was applied on the STAR-generated gene counts, such that both library size and gene length were taken into account (**Additional File 5**). Salmon v1.9.0 (alignment-based mode) was used for TPM normalisation, starting from a STAR-generated bam file with transcript coordinates.

In order to compare *P. knowlesi* gene expression patterns to their *P. vivax* orthologues, *P. vivax* time course mRNA reads from Zhu et al. (2016) were downloaded from NCBI (GEO: GSE61252). Reads were aligned to the PvP01 reference genome (PlasmoDB v2 release 66) and counted using STAR v2.7.6, and TPM-normalised with Salmon v1.9.0 as described above for *P. knowlesi*. Although the Zhu et al. (2016) study contains two time courses for clinical isolates smru1 and smru2, only the time course of smru1 was included (7 time points), as the smru2 time course showed a higher degree of asynchrony (20).

To analyse the *P. falciparum* time course, mRNA reads of 25 time points from strain II3 (derived from 3D7; 2 replicates per time point) (90) were downloaded from NCBI (GSE144976), read trimmed (BBDuk v36.99), aligned to the 3D7 reference genome (PlasmoDB v3 release 67) using STAR v2.7.6b and counted using the R package Rsubread v2.16.1 (function featureCounts). The obtained counts were TPM-normalised using edgeR v4.0.16 (convertCounts).

To enable comparison of expression levels between time points (but not between genes), TPM-normalised counts of *P. knowlesi*, *P. vivax* and *P. falciparum* were first log_2_-transformed (using pseudocount 1) to stabilise transcript count variance, and subsequently Z-scored (centred to mean 0 with standard deviation 1). For all analyses except the identification of constitutively expressed genes, genes were filtered prior to Z-scoring: (i) genes with virtually no expression were removed (sum of TPM-log_2_-normalised expression levels of all time points <0.5), and (ii) genes with minimal temporal variation were removed, as Z-scoring would artificially inflate noise (log_2_ fold change between highest and lowest TPM-log_2_-normalised expression value <0.5, using a pseudocount of 1).

Heatmaps were created with the ComplexHeatmap package in R. To cluster genes by their temporal expression profiles across the IDC, Ward’s D2 hierarchical clustering with Euclidean distances was applied. Gene ontology (GO) enrichment analyses were performed with the TopGO package in R. Curated and computationally inferred GO annotations for the *P. knowlesi* strain H and *P. vivax* strain P01 were downloaded from PlasmoDB (release 67 and release 66, respectively). The TopGO weight01 algorithm was used to take the hierarchy of the GO terms into account, and the Fisher’s exact test was used to calculate raw p-values. A Bonferroni multiple testing error correction was applied to identify significantly enriched GO terms (adjusted p<0.05).

### Extraction of constitutively expressed genes

Constitutively expressed genes were identified based on low transcriptional variation over the 5 time points and high overall expression. Low variation was defined as a log _2_ fold change < 0.5 between the highest and lowest TPM-log_2_-normalised expression values and a standard deviation < 0.3 of TPM- log_2_-normalised expression levels of the 5 time points. High expression was defined as a summed TPM-log_2_-normalised expression across the five time points above the genome-wide median (24.99).

### Extraction of stage-specific biomarkers and their application to annotate parasite age in existing single-cell datasets

For each time point of the *P. knowlesi* time course, 20 biomarker genes were selected. Per time point, genes with a TPM-normalised expression that was >3-fold upregulated compared to every other time point were identified, from which the 20 genes with the highest TPM-normalised expression were selected as biomarkers (**Additional File 3**).

These biomarkers were then visualised on existing single-cell *P. knowlesi* transcriptome data, generated using a HIVE (28) or 10X Genomics (27) platform. The HIVE single-cell data were pre-processed and analysed as previously described (28). The 10X single-cell count matrix of Howick et al. (2019) was downloaded (https://zenodo.org/records/2843883: pk10xIDC_counts dataset) and analysed following the methods of Sauve et al. (2024). Per time point, the mean expression of the biomarkers selected was visualised using UMAP and feature plots (FeaturePlot) from the Seurat package in R (v5.3.0) (108). In addition, *P. berghei*-based annotation of blood stages – as described for *P. knowlesi* in the Malaria Cell Atlas (27) – was downloaded (https://zenodo.org/records/2843883: pk10xIDC dataset; stage_pred column in the phenotypic metadata was used as life stage predictor) and visualised on the UMAP plots to compare with the time point annotation from this study. Pseudotime trajectories were inferred with Slingshot (2.14.0) on UMAP embeddings to capture developmental progression. Average biomarker expression per time point was plotted along pseudotime using a LOESS fit to highlight expression dynamics.

### Comparison of *P. knowlesi* expression patterns to *P. vivax* and *P. falciparum* orthologues

In order to compare gene expression patterns between *P. knowlesi* and *P. vivax* (31) or *P. falciparum* (90), IDC time (hpi) was normalised to a 0-1 scale to compare between the different IDC lengths of the species. Since different relative time points were sampled per species, the discrete TPM-normalised expression levels were smoothed and interpolated such that the full time range could be compared. The *P. falciparum* dataset (25 time points) was first smoothed using Locally Weighted Scatterplot Smoothing (LOWESS), in which the 5 closest time points were taken into account (Python statsmodels.nonparametric.smoothers_lowess.lowess). The *P. knowlesi* and *P. vivax* time courses were not smoothed, as they were too sparse (5 and 7 time points, respectively) for LOWESS. Next, to interpolate data points in between the first and last sampled time point, a Piecewise Cubic Hermite Interpolating Polynomial (PCHIP, PchipInterpolator from Python scipy.interpolate) was applied on the smoothed *P. falciparum* data and non-smoothed *P. knowlesi* and *P. vivax* data, with a step size of 0.005 relative time units. Interpolated data were then treated in the same way as the discrete TPM-normalised data: a log_2_ transformation was applied, and the same genes that were filtered out in the discrete data were removed. Finally, a Z-scoring was applied to enable comparison of relative temporal expression patterns between time points within genes, and across species.

To identify orthologous gene pairs, a protein BLAST from *P. vivax* to *P. knowlesi*, or from *P. falciparum* to *P. knowlesi* was carried out under the default settings. BLAST-resulting protein pairs with a % identity >=45% and with >=50% of the query protein aligned to its hit protein, were considered orthologues. For ApiAP2 transcription factors, *ApiAP2* genes with shared PlasmoDB gene names, a PlasmoDB OrthoMCL synteny hit or an indicated orthology in Jeninga et al. (2019) were considered orthologues as well, in case there was no hit among the BLAST-based orthologue list. To assess similarity of temporal expression dynamics between orthologous *P. vivax* - *P. knowlesi* gene pairs, dynamic time warping (DTW) was applied on the normalised interpolated expression levels (TPM-normalised, log_2_-transformed, Z-scored) over the time period that was sampled in both species (relative time 0.185 – 1) allowing for a temporal shift of 0.2 relative time units (R package dtw, Sakoe-Chiba window type). To determine a DTW score threshold to distinguish similar and dissimilar orthologue pairs, a baseline DTW score for dissimilar genes was generated: the transcript levels of every *P. vivax* gene were randomly shuffled to destroy the temporal structure and then matched to its *P. knowlesi* orthologue(s). **Figure 9** shows how the DTW score, baseline DTW score and time difference between peak expression levels of *P. vivax* and *P. knowlesi* (taking into account circularity of the IDC) were used to determine which orthologous gene pairs have a similar or dissimilar expression pattern over the IDC. **Additional File 4** lists the similarity classification and underlying statistics for all *P. vivax* – *P. knowlesi* orthologues.

**Figure 9.**
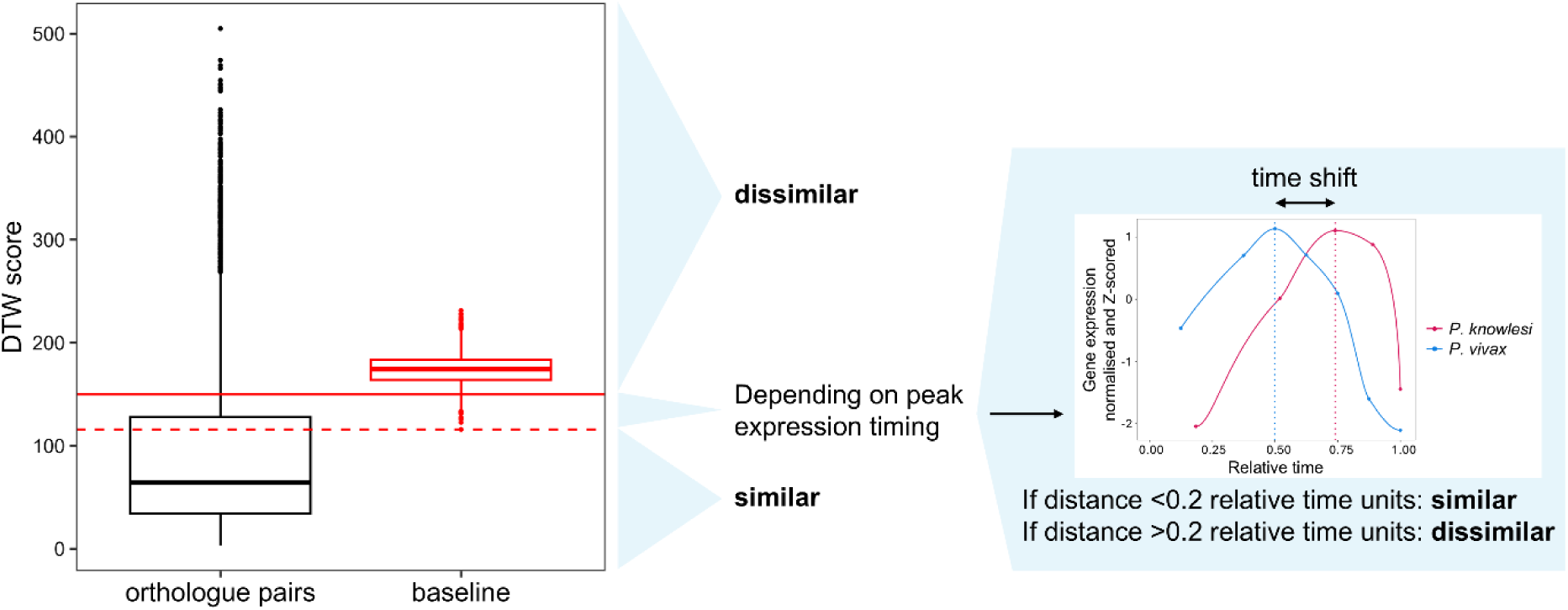
Classification of *P. vivax – P. knowlesi* orthologue pairs into similar or dissimilar IDC expression patterns. Box plots show the distribution of dynamic time warping (DTW) scores of orthologous gene pairs (black) and of the baseline DTW score distribution (red; random shuffling of the *P. vivax* expression levels per gene). The solid red line indicates the 5% quantile of the baseline DTW scores (149.7); orthologous gene pairs with DTW values above this line are considered to have a dissimilar expression pattern. The dotted red line indicates the minimum value of the baseline DTW scores (115.8); orthologous gene pairs with DTW values below this line are considered to have a similar expression pattern. When the DTW value is in between the 5% quantile and the minimum value of the baseline, orthologous gene pairs whose peak expression time points are <0.2 relative time units apart from each other (‘time shift’), are considered similar, if >0.2 relative time units apart from each other, dissimilar. Circularity of the IDC is taken into account when calculating time shift, meaning that relative times 0 and 1 have a time shift of 0.

An R shiny web tool was built in which temporal expression patterns of *P. vivax* - *P. knowlesi* orthologous gene pairs, as well as any *P. knowlesi* or *P. vivax* gene(s) of choice, can be visualised (https://interactive.itg.be/app/mal-pk-pv-expression-viewer). The underlying R code and datasets can be found on https://github.com/kdemeulenaere/Pk-Pv_Expression_Viewer.

### Whole-genome sequencing (WGS)

To verify presence of each *DBP* gene (*DBPα*, *DBPβ*, *DBPγ*) in our Pk A1-H.1 line, and to assess integrity of the *AP2-G* and *GDV1* regions, DNA was isolated from a blood sample collected in RNAprotect as part of the same experiment in which RNA samples for transcriptional analysis were collected. DNA was extracted using the QIAamp DNA Mini kit (Qiagen, #56304), following the manufacturer’s instructions. Elution in 50 µL of water yielded DNA at 5.0 ng/µL.

80 ng of DNA was sent to Novogene (Munich, Germany) for library preparation and sequencing (‘Microbial WGS’, including 10 cycles of PCR amplification). The 150 base pair paired-end Illumina sequencing on a NovaSeq X Plus system generated 3.5 million read pairs, resulting in an average sequencing depth of 42. The obtained fastq files are publicly available in the European Nucleotide Archive (ENA; https://www.ebi.ac.uk/ena/) under Study PRJEB83831.

Reads were aligned to the *P. knowlesi* strain H reference genome (PlasmoDB v2 release 66) using BWA-MEM (v0.7.19). 86.5% of all read pairs per sample mapped to a unique location on the *P. knowlesi* genome, and 18.5% of these mapped reads were marked as duplicates by Picard MarkDuplicates (v3.3.0). Aligned read coverage can be visualised in IGV using the IGV session file available on Zenodo (https://doi.org/10.5281/zenodo.18608274). Coverage tracks displayed in IGV were created with deepTools (v3.5.5), and only non-duplicate primary alignments with a BWA-MEM mapping quality of 60 were included.

Variants were called with GATK (v4.6.2.0; HaploTypeCaller, GenotypeGVCFs), and variant filtration was done separately on a single-nucleotide polymorphism (SNP) and insertion/deletion (INDEL) vcf file using the GATK golden standard settings (SNP vcf: QD<2.0, QUAL<30.0, SOR>3.0, FS>60.0, MQ<40.0, MQRankSum<-12.5, ReadPosRankSum<-8.0; INDEL vcf: QD<2.0, QUAL<30.0, FS>200.0, ReadPosRankSum<-20.0).

## Additional material

**Additional File 1 (.pdf). Supplementary data.** Supplementary Figures S1-S4 and Supplementary Table S1.

**Additional File 2 (.xls). Stage-specific GO terms.** A gene ontology (GO) enrichment analysis was carried out on gene sets grouped by their peak expression time point (5, 14, 20, 24 or 27 hpi; ‘time point gene sets’), against a background of all *P. knowlesi* genes. Only genes with sufficient expression and temporal variability were included (filtering criteria detailed in Methods). **Tab 1** (‘GO terms + statistics’) shows significantly enriched GO terms (p<0.05, and Bonferroni adjusted p<0.05) and their statistics for the 5 stage-specific gene sets (grouped according to their peak expression time point), separated over the ontology categories Molecular Function, Biological Process and Cellular Compartment. Colours indicate GO terms that are shared between different time point groups. **Tab 2** (‘Summary per category’) summarises the change in significantly enriched GO terms over the 5 time point gene sets, per GO category. **Tab 3** (‘% of genes with a GO term’) indicates the percentage of genes that are annotated with a GO term per time point gene set.

**Additional File 3 (.xls). List of the 20 biomarker genes that were selected for each time point.** Per time point, TPM-normalised genes showing a >3-fold upregulation compared to every other time point were identified, from which the 20 highest expressed genes at that time point were selected as biomarkers.

**Additional File 4 (.xls). Similarity classification for all *P. vivax* – *P. knowlesi* orthologous gene pairs.** In addition to the similarity classification (similar or dissimilar), the table also displays the statistics underlying this classification per orthologue pair: time points of peak expression for the *P. knowlesi* and the *P. vivax* gene, time shift between the *P. knowlesi* and *P. vivax* peak expression time point, and dynamic time warping score.

**Additional File 5 (.xls). TPM-normalised gene counts of the 5 sampled time points collected for *P. knowlesi* A1-H.1.** Hpi: hours post-invasion.

## Declarations

### Ethics approval and consent to participate

Not applicable.

### Consent for publication

Not applicable.

### Availability of data and materials

The Pk A1-H.1 transcriptome and whole-genome sequencing datasets are made publicly available in the European Nucleotide Archive (ENA; https://www.ebi.ac.uk/ena/) under study accession PRJEB83831, and mapped read coverage from these datasets can be visualised in Integrated Genome Viewer (IGV) using the IGV session file available on Zenodo (https://doi.org/10.5281/zenodo.18608274).

The interactive web tool for visualisation of *P. knowlesi* and *P. vivax* (orthologous) gene expression patterns is available at https://interactive.itg.be/app/mal-pk-pv-expression-viewer, the underlying R code and datasets can be found on Github (https://github.com/kdemeulenaere/Pk-Pv_Expression_Viewer) and Zenodo (https://doi.org/10.5281/zenodo.18599142).

Published datasets analysed in this study, are a *P. vivax* time course available in NCBI Gene Expression Omnibus (https://www.ncbi.nlm.nih.gov/geo/) under accession GSE61252 (31), a *P. falciparum* time course available in Gene Expression Omnibus under accession GSE144976 (90), and Pk A1-H.1 single-cell RNA-seq datasets available in Gene Expression Omnibus under accession GSE271911 (HIVE data) (28) and in ENA under accession ERP110344 (10X data) (27).

## Authors’ contributions

ARU and KDM conceptualised the study. ARU secured funding and provided overall supervision of the project. EK provided the Pk A1-H.1 lab strain and guidance on culturing, DDD and KDM carried out *P. knowlesi* culturing and time course sample collection, while ES contributed to the GuHCl procedure. KDM processed and analysed the sequencing data and generated the figures, with guidance from ARU and AC. KDM developed the web tool to visualise *P. knowlesi* and *P. vivax* temporal IDC expression. The *P. knowlesi* A1-H.1 HIVE single-cell data used in this paper was generated by ES (28); PM analysed the single-cell data shown in this paper and generated corresponding figures. The original draft of the paper was written by KDM and ARU; EK and AC were major contributors in editing the manuscript. All authors read and approved the final manuscript.

## Acknowledgements

We thank César Martinez for providing advice on LOWESS smoothing and interpolation, and for providing the processed *P. falciparum* transcriptome using data from Subudhi et al. (2020). We are grateful to Pieter Moris for his help with deploying the web tool on the server.

