## Additional File 1 for "A *Plasmodium knowlesi* A1-H.1 transcriptome time course focusing on the late asexual blood stages"

### Supplementary data

#### Supplementary Figures

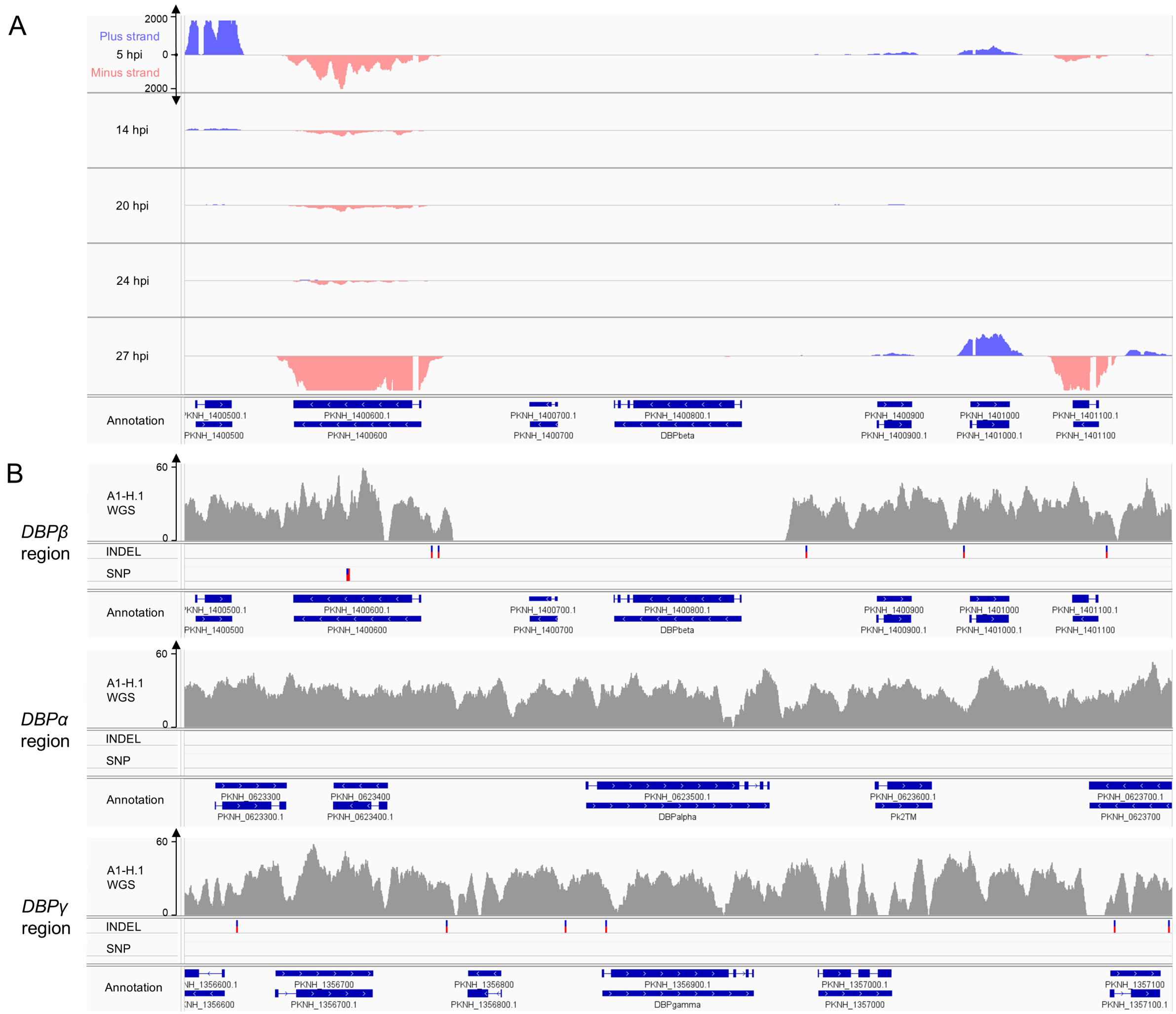


**Supplementary Figure S1. Integrated Genome Viewer images of *DBP* read coverage.** Coverage tracks were created with deepTools (v3.5.5). **(A)** mRNA-seq read coverage of the *DBPβ* gene and surrounding region for all 5 Pk A1-H.1 time point samples in this study. No *DBPβ* gene expression can be observed. Reads mapping to the plus strand are shown in blue, reads mapping to the minus strand in pink. Read depth for both strands is shown in a range of 0–2000, as illustrated on the y-axis of the 5 hpi sample (similar for all other time points). Only primary alignments with a STAR mapping quality of 255 are shown. The genome annotation (strain H) is shown at the bottom. For each gene, the full-length gene (no UTR's annotated for strain H), and the gene with exons (thick boxes) and introns (connecting lines) is displayed. Arrows indicate the strand on which the gene is located. **(B)** Whole-genome sequencing (WGS) read coverage of the *DBPα*, *DBPβ* and *DBPγ* gene and their surrounding region for Pk A1-H.1. Single-nucleotide polymorphisms (SNPs) and insertions/deletions (INDELs) are visible as vertical lines: blue line = homozygous mutation; blue-and-red line = heterozygous mutation (low-confidence since haploid). Read depth is shown in a range of 0–60. Only primary, non-duplicate alignments with a BWA-MEM mapping quality of 60 are shown. Annotation is as described for panel A.

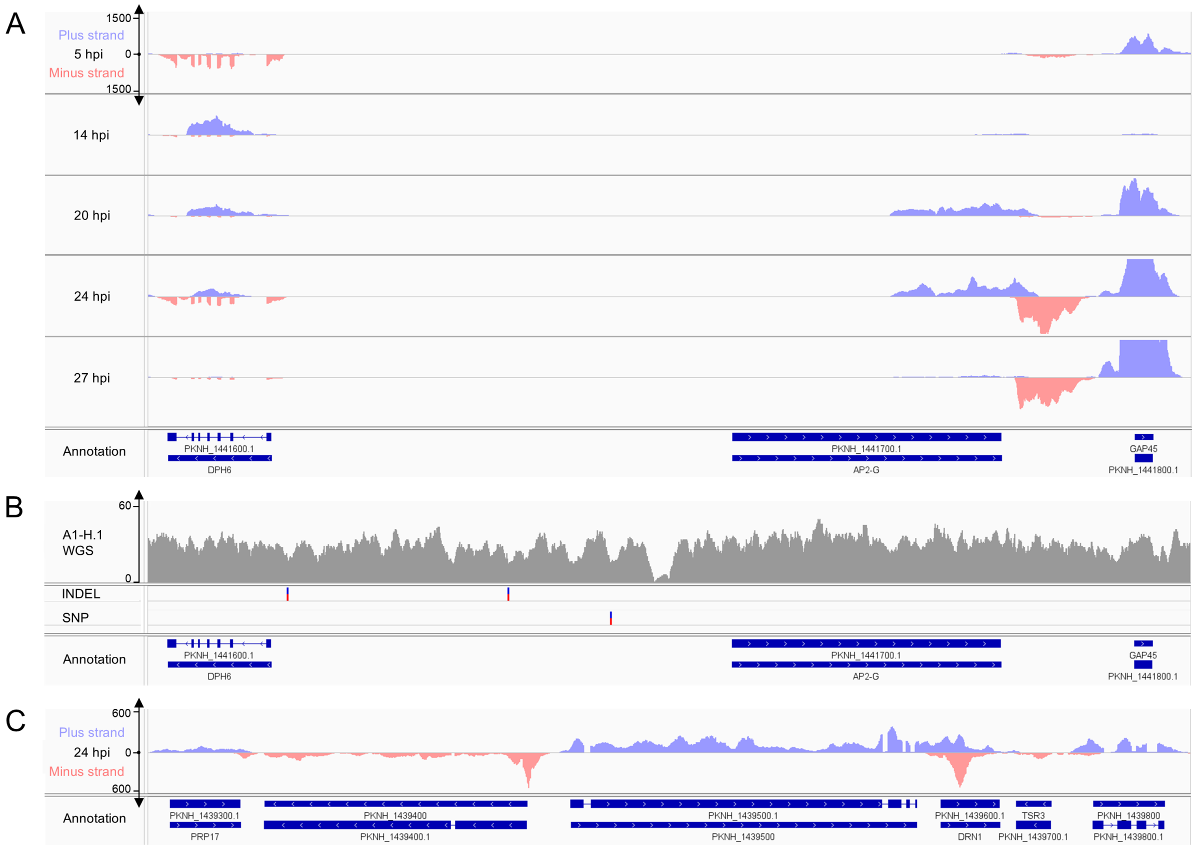


**Supplementary Figure S2. Integrated Genome Viewer images of *AP2-G* read coverage.** Coverage tracks were created with deepTools (v3.5.5). **(A)** mRNA-seq read coverage of the *AP2-G* gene and surrounding region for all 5 Pk A1-H.1 time point samples in this study. Reads mapping to the plus strand are shown in blue, reads mapping to the minus strand in pink. Read depth for both strands is shown in a range of 0–1500, as illustrated on the y-axis of the 5 hpi sample (similar for all other time points). Only primary alignments with a STAR mapping quality of 255 are shown. The genome annotation (strain H) is shown at the bottom. For each gene, the full-length gene (no UTR's annotated for strain H), and the gene with exons (thick boxes) and introns (connecting lines) is displayed. Arrows indicate the strand on which the gene is located. **(B)** Whole-genome sequencing (WGS) read coverage of the *AP2-G* gene and surrounding region for Pk A1-H.1. Single-nucleotide polymorphisms (SNPs) and insertions/deletions (INDELs) are visible as vertical lines: blue line = homozygous mutation; blue-and-red line = heterozygous mutation (low-confidence since haploid). Read depth is shown in a range of 0–60. Only primary, non-duplicate alignments with a BWA-MEM mapping quality of 60 are shown. Annotation is as described for panel A. **(C)** mRNA-seq read coverage of a chromosome 14 region containing two genes with a similar length as *AP2-G* (same genome region size as in panel A), shown for the 24 hpi sample. No strong 5' RNA degradation (absence of transcripts in this region) can be observed. Reads mapping to the plus strand are shown in blue, reads mapping to the minus strand in pink. Read depth for both strands is shown in a range of 0–600. Only primary alignments with a STAR mapping quality of 255 are shown. Annotation is as described for panel A.

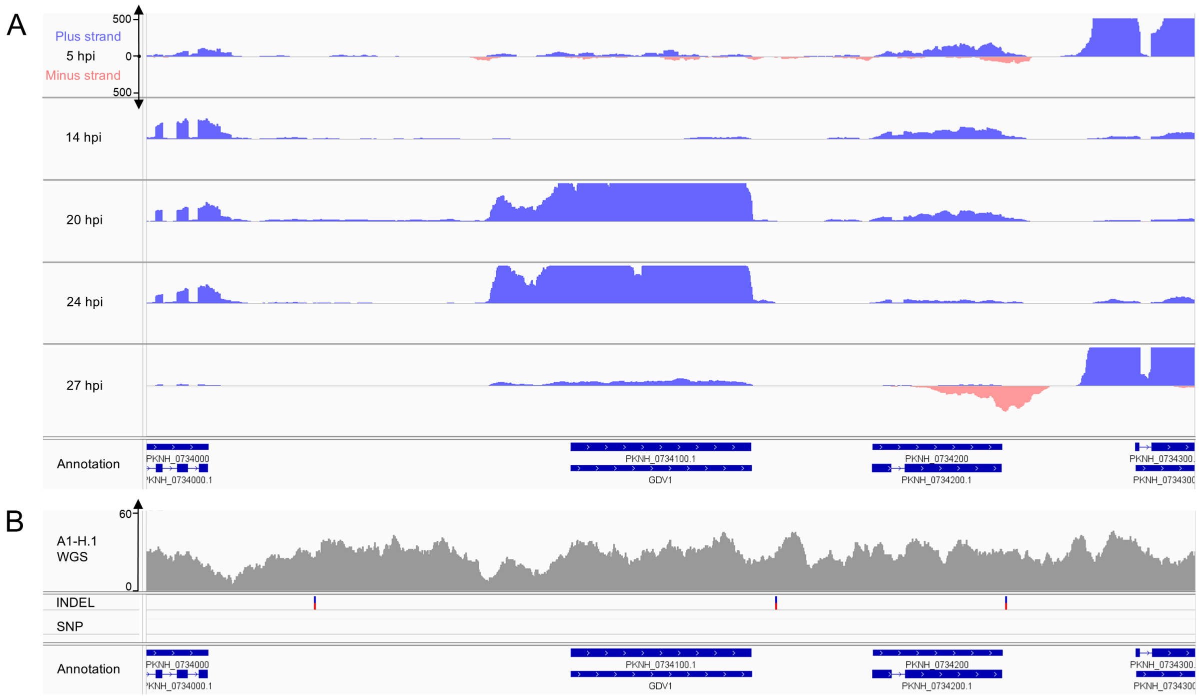


**Supplementary Figure S3. Integrated Genome Viewer screenshots of *GDV1* read coverage.** Coverage tracks were created with deepTools (v3.5.5). **(A)** mRNA-seq read coverage of the *GDV1* gene and surrounding region for all 5 Pk A1-H.1 time point samples in this study. Reads mapping to the plus strand are shown in blue, reads mapping to the minus strand in pink. Read depth for both strands is shown in a range of 0–500, as illustrated on the y-axis of the 5 hpi sample (similar for all other time points). Only primary alignments with a STAR mapping quality of 255 are shown. The genome annotation (strain H) is shown at the bottom. For each gene, the full-length gene (no UTR's annotated for strain H), and the gene with exons (thick boxes) and introns (connecting lines) is displayed. Arrows indicate the strand on which the gene is located. **(B)** Whole-genome sequencing (WGS) read coverage of the *GDV1* gene and surrounding region for Pk A1-H.1. Single-nucleotide polymorphisms (SNPs) and insertions/deletions (INDELs) are visible as vertical lines: blue line = homozygous mutation; blue-and-red line = heterozygous mutation (low-confidence since haploid). Read depth is shown in a range of 0–60. Only primary, non-duplicate alignments with a BWA-MEM mapping quality of 60 are shown. Annotation is as described for panel A.

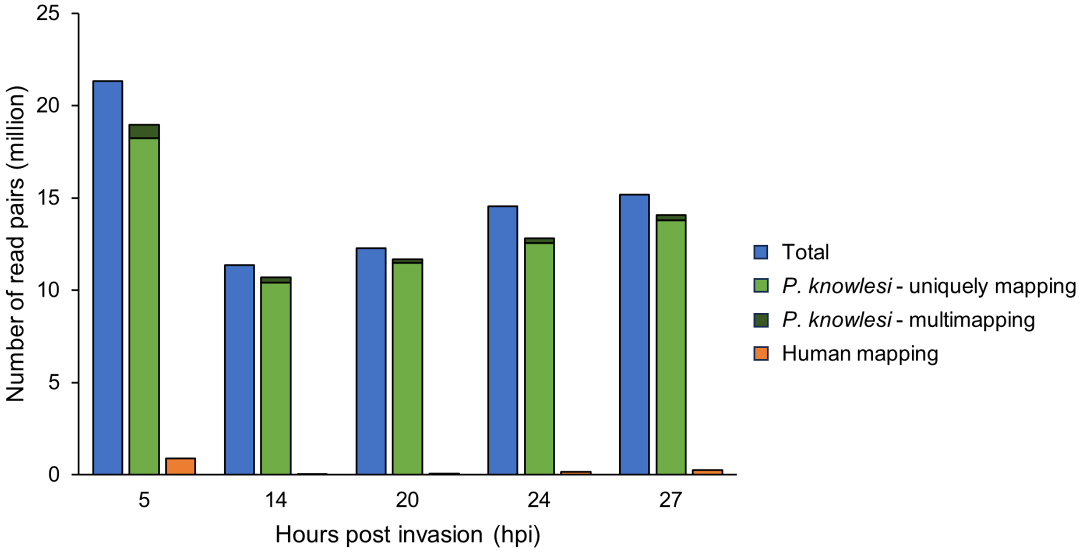


**Supplementary Figure S4. Overview of the number of obtained read pairs per time point**. 'Total' refers to all generated read pairs (mapping to the *P. knowlesi* or human genome, and unmapped reads). Reads mapping to the *P. knowlesi* genome H are split in uniquely mapping reads that map only to one location in the genome, and multimapping reads that map to multiple locations in the *P. knowlesi* genome. For reads mapping to the human genome, uniquely mapping and multimapping reads are counted together.

### Supplementary Tables

**Supplementary Table S1. List of the 10 highest expressed genes per time point.** Genes were TPM-normalised, and only nuclear-encoded genes were included (excluding mitochondrion- or apicoplast-encoded genes). Colours indicate genes that are shared between different time points.

| | **Gene ID** | **Product description** | **Gene name** |
|---|---|---|---|
| **Rings - 5 hpi** | | | |
| 1 | PKNH_0418600 | early transcribed membrane protein | ETRAMP |
| 2 | PKNH_1100900 | conserved Plasmodium protein, unknown function | N/A |
| 3 | PKNH_0835700 | 40S ribosomal protein S23, putative | N/A |
| 4 | PKNH_1355500 | ribosomal protein S27a, putative | N/A |
| 5 | PKNH_0734900 | Plasmodium exported protein, unknown function | N/A |
| 6 | PKNH_1325700 | knob-associated histidine-rich protein, putative | KAHRP |
| 7 | PKNH_1325800 | Plasmodium exported protein, unknown function | N/A |
| 8 | PKNH_0835500 | 60S ribosomal protein L7, putative | N/A |
| 9 | PKNH_1016300 | 60S ribosomal protein L12, putative | N/A |
| 10 | PKNH_0315000 | 60S ribosomal protein L11a, putative | N/A |
| **Mid-trophozoites - 14 hpi** | | | |
| 1 | PKNH_0515700 | early transcribed membrane protein | ETRAMP |
| 2 | PKNH_1218700 | glyceraldehyde-3-phosphate dehydrogenase, putative | GAPDH |
| 3 | PKNH_0418600 | early transcribed membrane protein | ETRAMP |
| 4 | PKNH_0614300 | adenosine deaminase, putative | ADA |
| 5 | PKNH_1114400 | elongation factor 1-alpha, putative | N/A |
| 6 | PKNH_0818800 | peptidyl-prolyl cis-trans isomerase, putative | CYP19A |
| 7 | PKNH_1114500 | elongation factor 1-alpha | N/A |
| 8 | PKNH_1426100 | DNA/RNA-binding protein Alba 1, putative | ALBA1 |
| 9 | PKNH_1259400 | 40S ribosomal protein S6, putative | N/A |
| 10 | PKNH_1349800 | 40S ribosomal protein S8e, putative | N/A |
| **Late trophozoites - 20 hpi** | | | |
| 1 | PKNH_1448000 | merozoite surface protein 9 | MSP9 |
| 2 | PKNH_0902700 | histone H2B, putative | H2B |
| 3 | PKNH_1457400 | merozoite surface protein 3, putative | N/A |
| 4 | PKNH_0819900 | histone H2A.Z, putative | H2A.Z |
| 5 | PKNH_1266000 | merozoite surface protein 7D | MSP7D |
| 6 | PKNH_1312700 | heat shock protein 70, putative | HSP70 |
| 7 | PKNH_1258400 | phosphoethanolamine N-methyltransferase | PMT |
| 8 | PKNH_1132600 | histone H2A, putative | H2A |
| 9 | PKNH_0715900 | endoplasmic reticulum chaperone BiP, putative | BIP |
| 10 | PKNH_0413400 | cysteine protease, putative | N/A |
| **Mid-schizonts - 24 hpi** | | | |
| 1 | PKNH_1448000 | merozoite surface protein 9 | MSP9 |
| 2 | PKNH_1132600 | histone H2A, putative | H2A |
| 3 | PKNH_0902700 | histone H2B, putative | H2B |
| 4 | PKNH_0611700 | peroxiredoxin, putative | nPrx |

| | | | |
|---|---|---|---|
| 5 | PKNH_1457400 | merozoite surface protein 3, putative | N/A |
| 6 | PKNH_1132500 | histone H3 variant, putative | H3.3 |
| 7 | PKNH_0703100 | high molecular weight rhoptry protein 3, putative | RhopH3 |
| 8 | PKNH_1347900 | rhoptry-associated protein 1 | RAP1 |
| 9 | PKNH_0819900 | histone H2A.Z, putative | H2A.Z |
| 10 | PKNH_0946200 | conserved Plasmodium protein, unknown function | N/A |
| **Late schizonts - 27 hpi** | | | |
| 1 | PKNH_0418600 | early transcribed membrane protein | ETRAMP |
| 2 | PKNH_0205200 | ELM2 domain-containing protein, putative | N/A |
| 3 | PKNH_1401700 | Plasmodium exported protein, unknown function | N/A |
| 4 | PKNH_0100400 | Plasmodium exported protein, unknown function | N/A |
| 5 | PKNH_0321400 | early transcribed membrane protein | ETRAMP |
| 6 | PKNH_0919300 | exported protein 1, putative | EXP1 |
| 7 | PKNH_1247400 | Plasmodium exported protein, unknown function | N/A |
| 8 | PKNH_1121500 | choline/ethanolaminephosphotransferase, putative | CEPT |
| 9 | PKNH_0802400 | inner membrane complex protein 1c, putative | IMC1c |
| 10 | PKNH_1465800 | actin I, putative | ACT1 |
